# Adding a new dimension: Multi-level structure and organization of mixed-species *Pseudomonas aeruginosa* and *Staphylococcus aureus* biofilms in a 4-D wound microenvironment

**DOI:** 10.1101/2022.05.14.491929

**Authors:** Radhika Dhekane, Shreeya Mhade, Karishma S Kaushik

## Abstract

Biofilms in wounds typically consist of aggregates of bacteria, most often *Pseudomonas aeruginosa* and *Staphylococcus aureus*, in close association with each other and the host microenvironment. Given this, the interplay across host and microbial elements, including the biochemical and nutrient profile of the microenvironment, likely influences the structure and organization of wound biofilms. While clinical studies, *in vivo* and *ex vivo* model systems have provided insights into the distribution of *P. aeruginosa* and *S. aureus* in wounds, they are limited in their ability to provide a detailed characterization of biofilm structure and organization across the host-microbial interface. On the other hand, biomimetic *in vitro* systems, such as host cell surfaces and simulant media conditions, albeit reductionist, have been shown to support the co-existence of *P. aeruginosa* and *S. aureus* biofilms, with species-dependent localization patterns and interspecies interactions. Therefore, composite *in vitro* models that bring together key features of the wound microenvironment could provide unprecedented insights into the structure and organization of mixed-species biofilms. We have built a four-dimensional (4-D) wound microenvironment consisting of a 3-D host cell scaffold of co-cultured human epidermal keratinocytes and dermal fibroblasts, and an *in vitro* wound milieu (IVWM); the IVWM provides the fourth dimension that represents the biochemical and nutrient profile of the wound infection state. We leveraged this composite 4-D wound microenvironment to probe the structure of mixed-species *P. aeruginosa* and *S. aureus* biofilms across multiple levels of organization such as aggregate dimensions and biomass thickness, species co-localization and organization within the biomass, overall biomass composition and interspecies interactions. In doing so, the composite 4-D wound microenvironment platform provides multi-level insights into the structure of mixed-species biofilms, which we incorporate into the current understanding of *P. aeruginosa* and *S. aureus* organization in the wound bed.

## INTRODUCTION

Biofilms are multicellular microbial communities, implicated in chronic wound infections, where they fuel the persistent, non-healing wound state [1–4]. In chronic wounds, biofilms are seen as discrete, microbial aggregates distributed across, and in close association with, wound bed [5–11]. Under these conditions, biofilm aggregates interface with a range of microenvironmental factors including host cells, such as fibroblasts and keratinocytes, as well as matrix, biochemical and nutrient factors [12–15]. Further, biofilms in wounds often consist of more than one microbial species [3,9,10,16,17]; *Pseudomonas aeruginosa* and *Staphylococcus aureus* are the most common bacterial co-pathogens. Consequently, *P. aeruginosa* and *S. aureus* have been observed to co-exist in the wound bed, with distinct structural and organizational features. In biopsies of chronic wounds, *P. aeruginosa* and *S. aureus* aggregates exist in distinct regions, albeit separated by a few hundred micrometers [6]. On the other hand, in *in vivo* wound models the two species are seen to exist in close proximity, with bacterial clusters observed to overlap with each other [18–20]. While the presence of truly-mixed aggregates with both species might be debated, *P. aeruginosa* and *S. aureus* have been detected in chronic wound biofilm aggregates consisting of bacteria of different species and morphologies [1,9–11], including bacilli and cocci. Given this, it is very likely that during initial infection, or at some point in the progression of the infected wound state, *P. aeruginosa* and *S. aureus* exist in close proximity with each other, allowing for possible interspecies interactions [21,22]. Therefore, in the complex infection microenvironment, the interplay across host and microbial elements likely influences the structure and organization of mixed-species bacterial biofilms.

The vast majority of laboratory studies on wound-relevant biofilms employ two-dimensional (2-D) plastic surfaces and refined protein broths [23–27], in which biofilms are seen to grow as homogeneous dense mats, or mushroom-like microcolonies [28–30]. This is clearly different from conditions and observations in clinical wounds, and are therefore of limited relevance. While *in vivo* and *ex vivo* systems have provided more relevant insights into the dimensions and distribution of biofilm aggregates [6–8,20,31,32], they present technical and scientific challenges with respect to studying biofilm structure across the host-microbial interface. More recently, there has been a push towards developing engineered *in vitro* approaches that recapitulate key features of the wound infection state, and enable the study of wound biofilms in the context of the complex microenvironment [13,18,33,34]. These approaches include reconstructed *in vitro* systems, such as mixed-species biofilms on 3-D host cell surfaces and in simulant media conditions that mimic the wound milieu [13,18,33,35–37]. In these reconstructed systems, biofilm aggregates have been shown to display species-dependent localization patterns and characteristic interspecies interactions [13,18,33], underscoring their role as relevant and tractable systems to study the structure and organization of mixed-species biofilms.

To study the structure and organization of mixed-species biofilms under conditions that mimic the infection state, we have built a four-dimensional (4-D) wound microenvironment consisting of a reconstructed host cell surface and an *in vitro* wound milieu (IVWM). The 3-D host cell scaffold consists of human epidermal keratinocytes (HaCaT) and human dermal fibroblasts (HDFa), co-cultured and fixed to serve as the substratum for biofilm formation. In our previous work, we developed and evaluated an IVWM that recapitulates the composition of clinical wound fluid, consisting of serum along with matrix elements such as collagen, fibrinogen, and fibronectin, and biochemical factors such as lactic acid and lactoferrin [33]. In doing so, the IVWM provides an additional dimension that represents the biochemical and nutrient profile of the wound infection state. Using confocal fluorescence microscopy and quantitative image analysis, we leveraged this recapitulated 4-D microenvironment to characterize structure of mixed-species *P. aeruginosa* and *S. aureus* biofilms across multiple levels of organization, such as aggregate dimensions and biomass thickness, species colocalization and organization within the biomass, overall biomass composition and interspecies interactions. Taken together, our findings provide multi-level insights into mixed-species *P. aeruginosa* and *S. aureus* biofilms under wound-relevant conditions, which we incorporate into the current understanding of *P. aeruginosa* and *S. aureus* organization in the wound bed.

## MATERIALS AND METHODS

### Bacterial strains and growth conditions

*Pseudomonas aeruginosa* (PAO1-pUCP18::mCherry, Amp/Carb^R^) and *Staphylococcus* R *aureus* (Strain AH13-pAH13::GFP_uvr_, Erm^R^) were gifted by Dr. Derek Fleming and Dr. Kendra Rumbaugh (Texas Tech University Health Science Center, Lubbock, Texas, USA) [38,39]. For all experiments, *P. aeruginosa* was grown in Luria Bertani (LB) medium (broth: Sigma, USA, L3022; agar: SRL, India, 474236) containing 100 *μ*g/mL ampicillin (HiMedia, India, TC021), and *S. aureus* was grown in LB medium containing 10 *μ*g/mL erythromycin (Himedia, India, TC024). *P. aeruginosa* and *S. aureus* were streaked on antibiotic-containing LB agar and incubated at 37°C. For overnight cultures, isolated colonies from streaked agar plates were grown in antibiotic-containing LB broth at 37°C under shaking conditions for 18-20 hours.

### Host cell culture and maintenance

Primary human dermal fibroblasts (HDFa) were purchased from PromoCell (Germany, C-12302) and cultured in Fibroblast Growth Medium (FGM) (Cell Applications, USA 116-500; medium contains 2% fetal bovine serum (FBS)). The immortalized epidermal keratinocyte cell line (HaCaT) was a gift from Dr. Madhur Motwani (Linq Labs, Pune, India) and was cultured in Keratinocyte serum-free Growth Medium (KGM) (Cell Applications, USA, 131-500A) supplemented with 1% FBS (Gibco, Brazil, 10270106). The cells were grown in tissue-culture treated, 25 cm^2^ culture flasks (Tarsons, Korea, 950040) and maintained at 37°C in a 5% CO_2_ humidified incubator. For all experiments, HDFa cells used had passage numbers <10, and HaCaT cells had passage numbers between 10-20.

### Preparation of the *in vitro* wound milieu (IVWM)

The *in vitro* wound milieu (IVWM) was prepared with FBS as the base component, with the addition of collagen, fibronectin, fibrinogen, lactic acid, and lactoferrin, as previously described [33]. Fibrinogen (Sigma, USA, F3879) was dissolved in filter-sterilized (0.2 *μ*m pore size, Cole-Parmer, India, WW-15945-52), pre-warmed saline (0.9% NaCl, Merck, India, 106404) to the concentration of 10 mg/mL and stored at −20°C. Fibronectin (Sigma, USA, F4759) was dissolved in autoclaved distilled water to the concentration of 1 mg/mL and stored at −20°C. Peptone water (0.1% w/v) was prepared by dissolving peptone (SRL, India, 95292) in 0.9% NaCl, and autoclaved and stored at 4°C. Lactoferrin (2 mg/mL) (Sigma, USA, L4040) was prepared in PBS (pH 7.2, Thermo Fisher Scientific, USA, 20012027), filter-sterilized and stored at 4°C. Commercial FBS (Gibco, Brazil, 10270106) was stored at 4°C. Rat tail collagen (50 *μ*g/mL in 0.02 M acetic acid) (Sigma, USA, 12220) was stored at 4°C. Lactic acid ≥85%; Sigma, USA, W261114) was stored at room temperature. Components stored at −20°C were thawed on ice prior to reconstitution of the IVWM. The IVWM was prepared by adding the above components to FBS (final concentration in IVWM, 70%) as follows (numbers indicate final concentrations): fibrinogen (300 *μ*g/mL), fibronectin (30 *μ*g/mL), rat-tail collagen (11.66 *μ*g/mL), lactoferrin (20 *μ*g/mL) and lactic acid (10.9 mM). Reconstituted IVWM was freshly prepared for all experiments.

### Preparation of host cell scaffolds

HaCaT and HDFa cells were grown in cell culture flasks up to 80-90% confluency. Cells were trypsinized with 700 *μ*L of 0.25% Trypsin-EDTA solution (Gibco, Canada, 25200056), followed by the addition of 700 *μ*L KGM with 1% FBS (for HaCaT) or FGM (for HDFa). To prepare co-cultured scaffolds of HDFa and HaCaT cells, 50 *μ*L of 1×10^5^ cells/mL (~5000 cells of HaCaT cells in serum-free KGM) and 50 *μ*L of 1×10^5^ cells/mL of HDFa cells (~5000 cells in FGM containing 2% FBS) were added to a single well of 96 - well flat clear bottom, tissue culture treated, black polystyrene microplate (Corning, USA, 3603), and grown in a 5% CO_2_ humidified incubator at 37°C for 72 hours [40]. Following this, the supernatant medium was removed, and the confluent cells were fixed in 50 *μ*L of 4% paraformaldehyde (PFA) (Sigma, USA, 158127) for 10 minutes at room temperature. Fixed cells were rinsed in 200 *μ*L sterile PBS four times, and stored in 200 *μ*L sterile PBS at 4°C until further use. Prior to experiments, fixed cell scaffolds were stained with 50 *μ*L of filter-sterilized 30 *μ*M 4’,6-Diamidino-2-Phenylindole, Dihydrochloride (DAPI) (Invitrogen, Germany, D1306) for 5 minutes at room temperature, followed by two rinses with 200 *μ*L PBS.

### Growth of biofilms in the composite 4-D wound microenvironment

Overnight cultures of *P. aeruginosa* and *S. aureus* (in LB media) were each diluted 2×10^7^ cells/mL in freshly-prepared IVWM (after addition of bacterial cultures the medium would contain less than 2% LB). For mixed-species biofilms, 50 *μ*L of *P. aeruginosa* and 50 *μ*L of *S. aureus* diluted in IVWM (~10^6^ cells each) were added to fixed HDFa-HaCaT host cell scaffolds. For single-species biofilms, 50 *μ*L of *P. aeruginosa* or *S. aureus* diluted in IVWM (~10^6^ cells) were added to fixed HaCaT+HDFa host cell scaffolds, followed by the addition of 50 *μ*L of IVWM to make a total volume of 100 *μ*L. The composite 4D microenvironment, including microbial cells, host cell scaffolds and the IVWM, was incubated at 37°C in a static incubator for 4, 8, 24 and 48 hours. Given the possible detrimental effects of laser exposure on bacterial growth, biofilm wells for different time points were seeded together and images for each time point were obtained from a different well.

### Confocal microscopy and image acquisition

At 4, 8, 24, and 48 hours, undisturbed *P. aeruginosa* and *S. aureus* biomass on fixed host cell scaffolds were imaged using confocal laser scanning microscopy (Leica, Germany, LASX TCS SP). Briefly, each well was imaged at the approximate center, with at least 3 biological replicates for each time point. To visualize GFP-labeled *S. aureus*, a 488 nm excitation filter and 497 nm to 542 nm emission filter was used. To visualize mCherry-labeled *P. aeruginosa*, a 561 nm excitation filter and 586 nm to 656 nm emission filters were used. To visualize DAPI-stained host cell nuclei, a 405 nm excitation filter and 414 nm to 485 nm emission filter was used. Biomass on fixed host cell scaffolds were imaged using a Z-stack of 100 *μ*m to 175 *μ*m thickness, with a 5 *μ*m step size. For imaging the host cell scaffolds alone (without biofilms), Z-stacks of 1 *μ*m step size and 60 *μ*m total thickness was acquired, with two fields of view per well.

### Image processing and analysis

Image analysis was done using the open-source image analysis tools, ImageJ and BiofilmQ v0.2.1., with Paraview v5.10.0 used for rendering [41–48].

To characterize the host cell scaffolds, a four-step process with BiofilmQ v0.2.1 was used [41,42]. This consisted of image preparation, image processing, calculation of features of the nuclei, and visualization. Briefly, each DAPI-stained image was subjected to separation using an intensity threshold filter to isolate individual nuclei. This calculated the total number of host cell nuclei in the visualized area. Each separated nucleus and its MATLAB parameter files were exported into a separate folder and processed independently for analysis. For image processing, each nucleus was subjected to semantic segmentation using the Otsu thresholding method [43]. Further, the images were subjected to de-noising by convolution (kernel size in pixels (xy,z)-5,3) and top-hat filter of size 25 vox (14.20 *μ*m) to remove background fluorescence. Next, object declumping by cube segmentation was used for dissection of each nuclei into smaller cubes (20 vox or 11.36 *μ*m). In addition, a small object removal filter was used to remove voxel clusters of size less than 5000 vox to refine the visualized area prior to parameter calculations. Local and global parameters in BiofilmQ v0.2.1 [41,42] were used for calculating height, shape-volume and base-area of the nuclei. During processing, visualizations were generated using the .vtk file output capability within BiofilmQ v0.2.1 [41,42] and rendered using the open-source tool Paraview v5.10.0 [44–46].

Aggregate sizes for *P. aeruginosa* and *S. aureus* in mixed-species and single-species biomass were estimated using ImageJ [47–49]. For this, the LIF files for each time point were imported into ImageJ [47,48] and maximum intensity Z-projections of relevant channels were obtained. Otsu’s automated thresholding method was applied to the Z-projection, and the areas of thresholded particles, representing aggregate sizes, were measured using the particle size analysis tool. Resulting aggregate size values were exported to a spreadsheet and grouped into the following aggregate size ranges: <5 *μ*m^2^, 5 −10 *μ*m^2^, 10 −20 *μ*m^2^, 20-50 *μ*m^2^, 50-100 *μ*m^2^, and >100 *μ*m^2^. For 8, 24 and 48 hour time points, the aggregate sizes were groups as <25 *μ*m ^2^, 25 −50 *μ*m ^2^, 50-100 *μ*m ^2^, 100-1000 *μ*m ^2^, 1000-10000 *μ*m^2^ and >10000 *μ*m^2^. Further, the percent count for each aggregate size range for each condition was calculated as the number of aggregates in a given size range/total number of aggregates x 100.

The thickness of *P. aeruginosa* biofilms under mixed-species and single-species conditions at 8, 24 and 48 hour time points was measured using ImageJ [47,48]. Briefly, the side view projection images of the Z-stacks (of the relevant channel) were imported in ImageJ [47,48]. For a given replicate, the biofilm boundary was defined manually using the polygon selection tool, and the thickness of the biofilm was measured across every 25 *μ*m on the X-axis. The mean of 23 measurements was considered the average thickness for the biofilm replicate. The final mean thickness at each time point represents the average thickness of three replicates.

To measure the co-localization of *P. aeruginosa* and *S. aureus* biomass with respect to each other, the 3D overlap parameter in BiofilmQ v0.2.1 [41,42] was used. This feature quantifies the volume overlap or co-localization of two fluorescent channels in a given space. A local parameter called ‘Correlation Local3dOverlap’ calculates the volume overlap in *μ*m^3^ of each object in channel 3 (GFP-*S. aureus*) with all segmented objects in channel 4 (mCherry-*P. aeruginosa*), as well as the volume overlap of each object in channel 4 with all segmented objects in channel 3. This provides a ‘Correlation LocalOverlapFraction’, which is obtained by dividing ‘Correlation_Local3dOverlap’ by the volume of the object. These values are then used to provide the user with a 3D overlap fraction for the entire channel or the Biofilm_OverlapFraction, defined as the sum of the Correlation LocalOverlapFraction for all objects in a specific fluorescence channel.

To quantify the spatial organization of *P. aeruginosa* and *S. aureus* aggregates in the mixed-species biomass, LIF files containing images of mixed-species biofilms for each time point were imported into ImageJ [47,48], following which the entire Z-stack for each relevant channel was thresholded using Otsu’s thresholding method. The mean gray value (MGV) (indicative of fluorescent intensities) of the thresholded areas for each Z-slice in a given Z-stack was calculated using the ‘measure stack’ tool in ImageJ [47,48]. MGVs were normalized to the highest MGV in a given dataset, and plotted as a distribution of biomass over Z-thickness.

Biomass composition of *P. aeruginosa* and *S. aureus* in mixed-species and single-species biofilms was calculated using MGV in ImageJ [47,48]. For this, LIF files for each time point were imported into ImageJ, and maximum intensity Z-projections of relevant channels were obtained, which was then subject to Otsu’s automated [43,50] thresholding method. MGVs were obtained for *P. aeruginosa* and *S. aureus* single-species biomass, as well as separately for *P. aeruginosa* and *S. aureus* in the mixed-species biomass, for each time point. The ratio of MGVs for *P. aeruginosa* to *S. aureus* in the mixed-species biomass, and for mixed-species to single-species biomass for *P. aeruginosa* to *S. aureus*, was obtained to analyze the composition of mixed-species biomass, and possible role of interspecies interactions across time points.

### Statistical analysis

Statistical analysis was performed using GraphPad Prism 6.01 for Windows, GraphPad Software, San Diego, California USA, www.graphpad.com [51]. A one-way or two-way ANOVA with Fisher’s LSD test for multiple comparisons was performed and a p-value of ≤0.05 was considered significant.

## RESULTS AND DISCUSSION

### A 4-D wound microenvironment with host cell scaffolds and an *in vitro* wound milieu (IVWM) to study the structure and organization of mixed-species biofilms

In contrast to standard laboratory studies on 2-D surfaces (**Figure 1A**), biofilms in wounds are observed as bacterial aggregates, attached to the surface of, and surrounded by, the wound bed [5,7,52]. The wound bed is made up of host cells such as epidermal keratinocytes and dermal fibroblasts [35,40], and is bathed in protein-rich wound fluid. Together, the cellular and matrix elements of the wound bed, along with the biochemical and nutrient profile of wound fluid, result in a characteristic wound milieu. Given this, biofilms in wounds form and exist in close approximation with the complex wound microenvironment.

**Figure 1:**
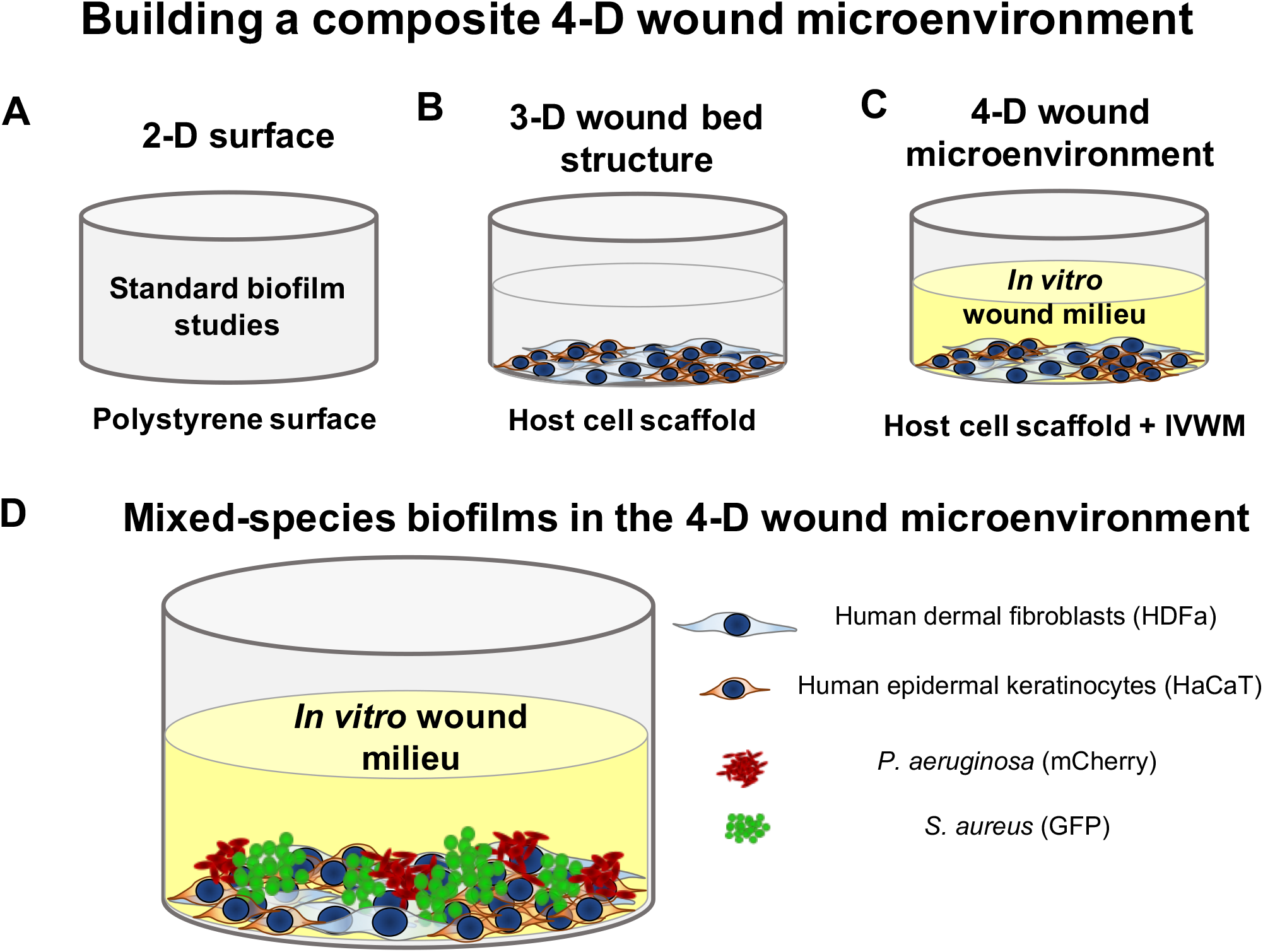
Building a 4-D wound microenvironment consisting of mixed-species *P. aeruginosa* and *S. aureus* biofilms on a co-cultured host cell scaffold of epidermal keratinocytes and dermal fibroblasts, in the presence of an *in vitro* wound milieu. **(A)** Standard biofilm studies typically use 2-D polystyrene surfaces of 96-well plates to grow and study biofilms. **(B)** Host cells, such as human dermal fibroblasts (HDFa) and epidermal keratinocytes (HaCaT), provide a 3-D surface that resembles the composition and organization of the wound bed. **(C)** The IVWM recapitulates the composition of clinical wound fluid, with matrix and biochemical factors, and provides the additional dimension that represents the microenvironmental profile of the infection state. **(D)** The composite 4-D wound microenvironment consists of mixed-species *P. aeruginosa* and *S. aureus* biofilms grown on the 3-D surface of semi-inert (fixed) HaCaT+HDFa host cell scaffold, in the presence of the IVWM.

To recapitulate the wound microenvironment, we developed an *in vitro* platform consisting of a 3-D reconstructed ‘wound bed’ and an *in vitro* wound milieu, which provides an additional dimension that represents the biochemical and nutrient profile of the infection state; this platform is collectively referred to as the 4-D wound microenvironment. The ‘wound bed’ was reconstructed using co-cultured HaCaT and HDFa cells, grown and fixed to form a confluent host cell scaffold **(Figure 1B)**. Based on our previous work [33], the *in vitro* wound milieu (IVWM) mimics the composition of clinical wound fluid, consisting of fetal bovine serum (FBS), with additional matrix and biochemical factors, such as collagen, fibrinogen, fibronectin, lactic acid and lactoferrin [33] **(Figure 1C)**. The IVWM supports the growth of mixed-species biofilms of *P. aeruginosa* and *S. aureus*, and recapitulates key features such as biomass formation and metabolic activity, and interspecies interactions [33].

In this study, the composite 4-D wound microenvironment was leveraged to study the structure and organization of *P. aeruginosa* and *S. aureus* biofilms under mixed-species conditions **(Figure 1D)**. To parse out the possible role of interspecies interactions, *P. aeruginosa* and *S. aureus* were also studied as single-species biofilms. It is important to note that while the presence of live host cells with features such as migration and proliferation, as opposed to fixed host cell scaffolds, would more closely mimic the wound bed, they present issues with respect to viability in the presence of bacterial growth [53]. This is particularly important in the context of studying biofilm structure and organization across time points, for which fixed host cell scaffolds provide a semi-inert but wound-relevant substratum.

### 3-D features of the co-cultured host cell scaffolds with HaCaT and HDFa cells

To reconstruct the ‘wound bed’, HaCaT and HDFa cells were co-cultured to form 3-D host cell scaffolds. Briefly, HaCaT and HDFa cells were mixed in a 1:1 ratio, and grown for 3 days, following which they were fixed and stained to visualize host cell nuclei. To characterize their features, fixed host cell scaffolds (without the addition of bacteria) were imaged using confocal microscopy. Images were analyzed with BiofilmQ v0.2.1 [41,42] using global parameters for number and height of the nuclei, and local parameters for the base area and shape-volume of the nuclei.

As seen in **Figure 2A** (inset images after fixation), the host cell scaffolds consisted of both HaCaT and HDFa cells, with characteristic co-culture arrangements. HaCaT cells were observed as closely-packed clusters, surrounded by sheaths of dermal fibroblasts (HDFa) [40], seen as large, flat, spindle-shaped cells, with elongated protrusions **(Figure 2A, inset)**. The co-cultured scaffolds showed an average of 140±16 cells (both HaCaT and HDFa) in a visualized area of 581.2 X 581.2 *μ*m, consisting of 88±16 HaCaT cells and 52±4 HDFa cells. This results in 13000±1500 cells in the well (culturable surface area of 0.32 X 10^8^ *μ*m^2^; n=2, two biological replicates with two fields of view per well, error represents SEM), which based on the average cell area for HaCaT and HDFa cells, 3000 *μ*m^2^ [54–57] and 4000 *μ*m^2^ [54–57] respectively, corresponds to a coverage of close to 100% of the well surface [58,59]. It is important to note that in the composite 4-D wound microenvironment, the IVWM would provide matrix elements (collagen, fibrinogen, fibronectin), which would be present between, and in close association, with the host cells.

**Figure 2:**
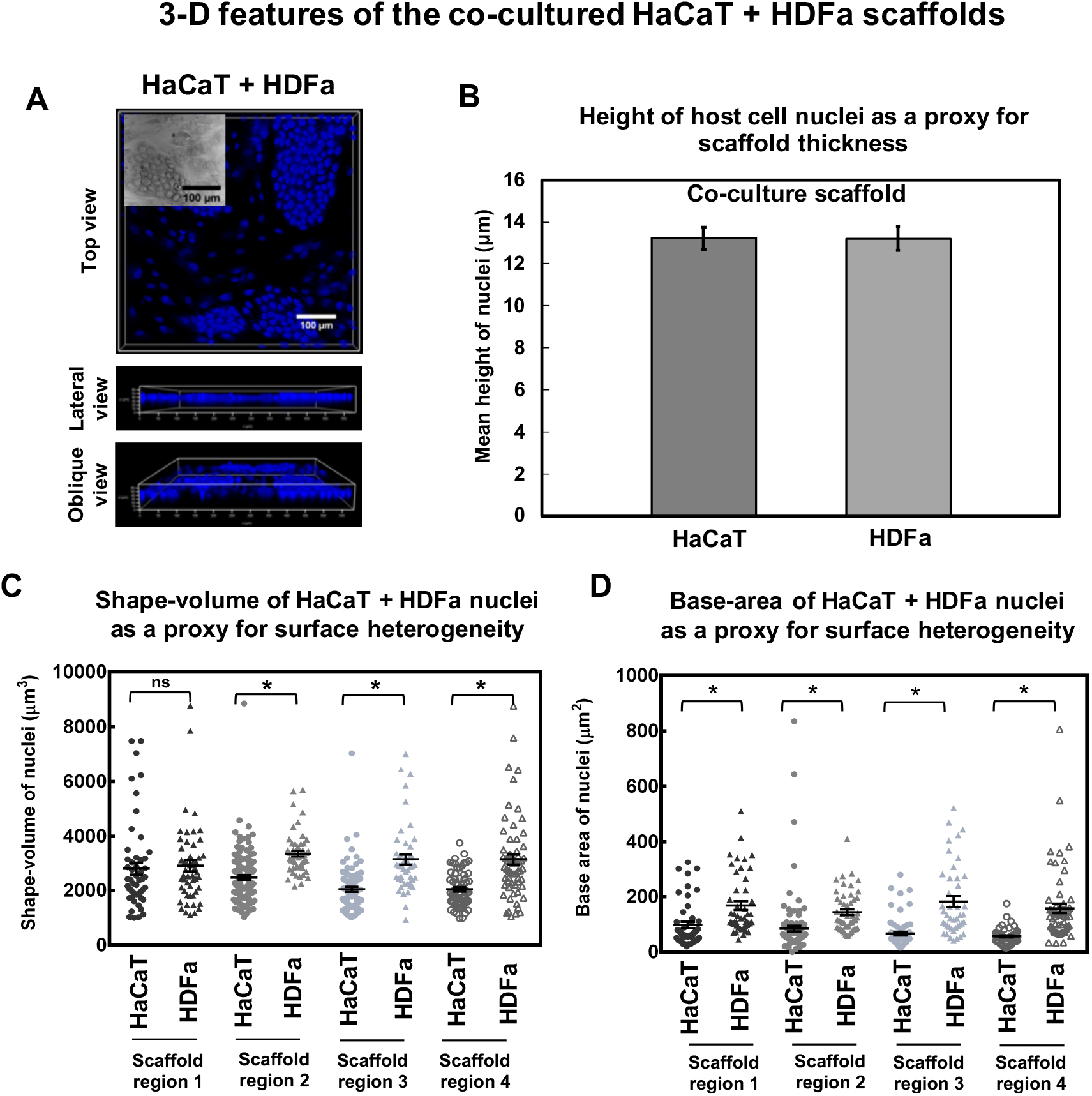
The co-cultured HaCaT+HDFa host cell scaffold provides a 3-D substrate that resembles the cellular and organizational structure of the wound bed. **(A)** Representative top, lateral and oblique views of the DAPI-stained host cell nuclei in co-cultured (fixed) HaCaT+HDFa scaffolds. **(B)** Mean height of the host cell nuclei (in *μ*m) in the HaCaT+HDFa scaffolds (n=2, error bars represent SEM), as a proxy for scaffold thickness. **(C)** Variation in shape-volume of host cell nuclei in HaCaT+HDFa across four scaffold replicates (n=2, error bars represent SEM), as a proxy for 3-D surface heterogeneity. **(D)** Variation in base area of host cell nuclei in HaCaT+HDFa across four scaffold replicates (n=2, error bars represent SEM), as a proxy for 3-D surface heterogeneity. Error bars represent SEM, n=2, indicating 2 biological replicates with 2 fields of view per well; ns indicates no significant difference, a p-value ≤0.05 was considered significant (*).

To characterize the 3-D features of the host cell scaffolds, we analyzed the height, shape-volume and base-area of the host cell nuclei (n=2, two biological replicates with two fields of view per well, error represents SEM). The average height of the HaCaT nuclei measured 13.2+0.5 *μ*m and that of the HDFa nuclei measured 13.2±0.6 *μ*m **(Figure 2B)**. It is important to note that the cytoplasmic layer surrounding the nucleus, typically 15-30 microns [60–62], would further add to the thickness of the substratum. The base-area of the nucleus represents the surface area occupied in the horizontal plane, and the shape-volume represents the volume occupied by the nucleus across both horizontal and vertical planes. Given that nuclear morphology varies with cell [63–65] shape and nuclear volume positively correlates with cell volume, base-area and shape-volume can serve as proxies for heterogeneity in cell shape and volume. As seen in **Figure 2C and D**, in the co-cultured HaCaT+HDFa scaffolds, host cell nuclei for each cell type showed variable distribution with respect to base-area and shape-volume. This heterogeneity is further underscored by the variation in base-area and shape-volume distribution across the two cell types **(Figures 2C and D)**.

Taken together, the co-cultured host cell scaffolds consist of a confluent layer of fixed HDFa and HaCaT cells, with 3-D features that recapitulate the cellular composition and organization of the wound bed.

### Mixed-species *P. aeruginosa* and *S. aureus* biofilms in the composite 4-D wound microenvironment

To study mixed-species biofilms in the composite 4-D microenvironment, *P. aeruginosa* and *S. aureus* were mixed in the IVWM, and seeded together (in a 1:1 ratio) on the 3-D surface of the HaCaT+HDFa fixed host cell scaffolds, followed by incubation at 37°C and imaging at 4, 8, 24 and 48 hours. For comparison with single-species biofilms, *P. aeruginosa* and *S. aureus* were mixed separately in the IVWM, and seeded alone.

As seen in **Figure 3A**, in the composite wound 4-D microenvironment, *P. aeruginosa* and *S. aureus* are observed to co-exist under mixed-species conditions, with the structure and organization of the biomass observed to vary across time points. At 4 hours, *P. aeruginosa* and *S. aureus* are seen as bacterial aggregates, in close approximation with, and dispersed across, the host cell scaffold (seen as DAPI-stained nuclei). Across the subsequent time points of 8, 24 and 48 hours, *P. aeruginosa* is observed to grow into dense, mat-like biofilms **(Figures 3B)**, whereas *S. aureus* retains its aggregate structure over time **(Figures 3C)**. Further, across 8, 24 and 48 hours, *S. aureus* aggregates are observed to be progressively enmeshed in the dense, mat-like growth of *P. aeruginosa* (**Figure 3A)**. Overall, this resembles the presence and growth of biofilms in the wound bed, observed as bacterial aggregates embedded in, and surrounded by, host cellular and matrix elements [6,7]. Further, *P. aeruginosa* aggregates in wounds are typically observed to be large and dense, extending across the wound surface area [6,66]. This is in contrast to *S. aureus* aggregates, which are most often seen as small and discrete clumps across the wound bed [6,7]. Notably, the visualized biomass consists of bacteria in aggregates or biofilms, as well as planktonic cells. In doing so, it resembles the structural complexity of biofilms in wounds, where bacteria can be seen as aggregates and dense clumps, as well as planktonic cells [9]. It is important to note that mixed-species biomass accumulation at later time points (likely an effect of *P. aeruginosa*) leads to the progressive destruction of the host cell scaffold, seen as diffusion of the nuclear stain through the visualized area **(Figure 3A and Supplementary Figure 1)**.

**Figure 3:**
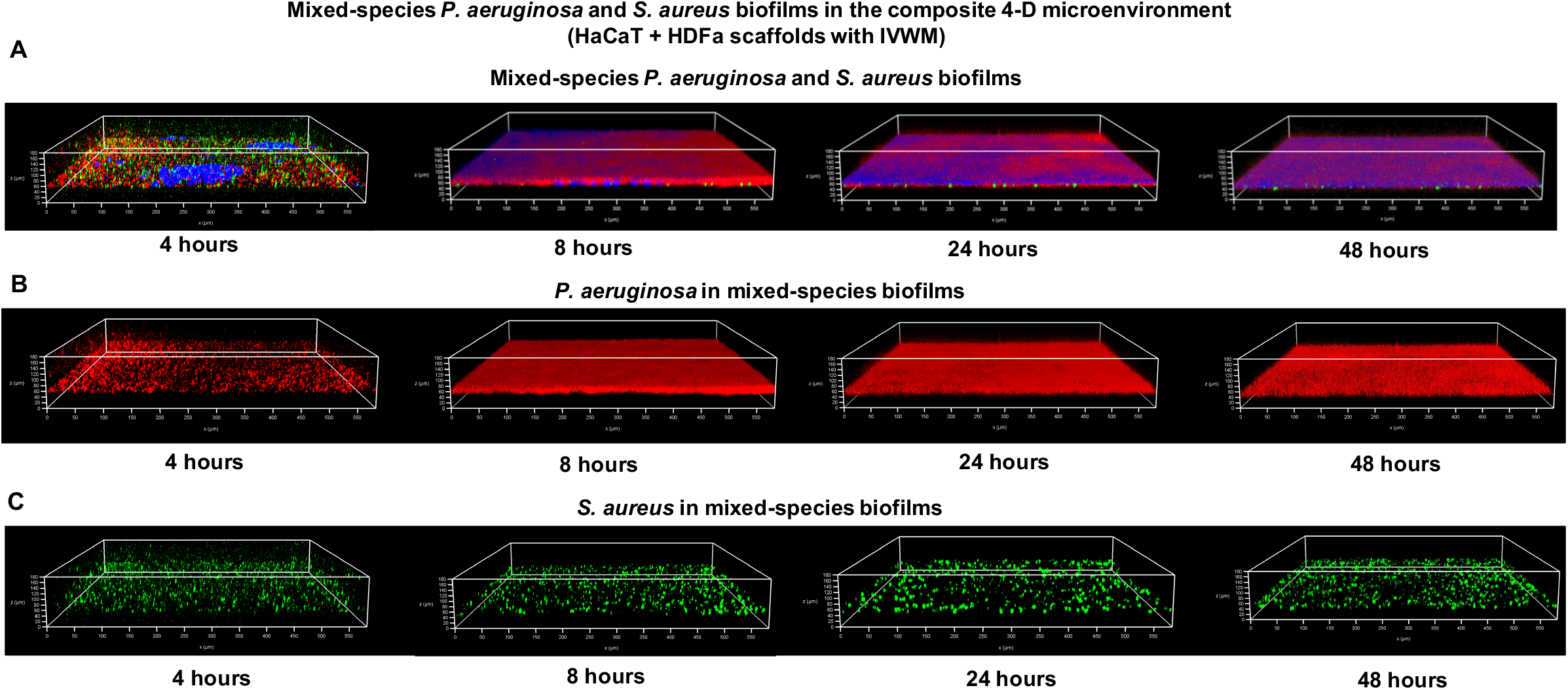
Mixed-species biomass of *P. aeruginosa* and *S. aureus* biofilms in the composite 4-D microenvironment show distinct structure and organization. *P. aeruginosa* (PAO1-pUCP18, mCherry) and *S. aureus* (Strain AH 133-pAH13, GFP) were seeded in a 1:1 ratio in IVWM on fixed HaCaT+HDFa scaffolds. Following incubation at 37°C, mixed-species biomass was imaged at 4, 8, 24 and 48 hours. **(A)** Representative images of mixed-species *P. aeruginosa* and *S. aureus* biomass in the composite 4-D microenvironment across time points. The blue stain represents the DAPI-stained host cell nuclei; the diffusion of the nuclear stain across subsequent time points is likely due to destruction of host cell structure with bacterial growth. **(B)** Representative images of *P. aeruginosa* in the mixed-species biomass (without the DAPI channel) in the composite 4-D microenvironment across time points. **(C)** Representative images of *S. aureus* in the mixed-species biomass (without the DAPI channel) in the composite 4-D microenvironment across time points.

Given this, the composite 4-D microenvironment platform provides a relevant and tractable system to study the structure of mixed-species biofilms across multiple levels of organization, such as aggregate dimensions and biomass thickness, species co-localization and organization within the biomass, and overall biomass composition. Further, using comparisons with single-species *P. aeruginosa* and *S. aureus* biofilms **(Supplementary Figure 2)**, the platform can be leveraged to parse out the roles of possible interspecies interactions.

### Aggregate dimensions and biomass thickness of *P. aeruginosa* and *S. aureus* mixed-species biofilms in the 4-D wound microenvironment

Aggregate sizes for *P. aeruginosa* (at 4 hours) and *S. aureus* (at 4, 8, 24 and 48 hours) in the mixed-species biofilms were measured using the particle size analysis tool in ImageJ [47–49], where the areas of thresholded particles correspond to aggregate size (measured in *μ*m^2^). It is important to note that while biofilm aggregates are 3-D structures, we measured the size of the aggregates in area (μm^2^), which is in accordance with previous studies [67]. Further, the biomass thickness of *P. aeruginosa* was measured at 8, 24 and 48 hours using ImageJ [47,48].

As seen in **Figure 4A**, early biofilms (visualized at 4 hours) of *P. aeruginosa* and *S. aureus* under mixed-species conditions, consist of a majority of single bacterial cells or small aggregates (<5 *μ*m^2^) that constitute 76% and 82% of the total biomass respectively. The next largest group of aggregates for both species are in the 5-10 *μ*m^2^ size range, representing 11% and 8% of the aggregate size distribution for *P. aeruginosa* and *S. aureus* respectively. This is followed by larger aggregates in the 10-20 *μ*m^2^ size range, that comprise ~5-6% of the total biomass. Taken together, the early mixed-species biofilm consists of ~70-80% of smaller aggregates of less than 10 *μ*m^2^ size, with larger aggregates (20-50 *μ*m^2^, 50-100 *μ*m^2^ and >100 *μ*m^2^ in size) comprising ~20-30% of the total biomass. The majority of single bacterial cells or small aggregates in early biofilm formation, corresponds to the well-studied model of biofilm development, where biofilms are typically seeded by single cells (planktonic) or small aggregates [28]. While the overall distribution of aggregate sizes (including the majority proportion of single cells and small aggregates) is similar for *P. aeruginosa* and *S. aureus* under mixed-species conditions, *S. aureus* biomass consisted of a significantly larger number of smaller aggregates (~82% in the <5 *μ*m^2^ size range), as compared with *P. aeruginosa* (~76%). On the other hand, *P. aeruginosa* biomass showed a shift towards larger aggregate sizes, such as in the 10-20 *μ*m^2^ and 20-50 *μ*m^2^ size ranges. Further, in the larger aggregate size range of >100 *μ*m^2^, *P. aeruginosa* biomass consists of ~1% of the aggregate populations, which is significantly higher than that for *S. aureus* (~0.1%) **(Figure 4A)**. It is important to note that while biofilms were seeded from bacterial cultures grown under planktonic conditions, overnight bacterial suspensions are known to contain a mixture of single cells and pre-formed aggregates [28,68–70]. This would contribute to a variation in the initial seeding of aggregates, and could thereby influence subsequent aggregate sizes, including the growth of a few large aggregates [28]. Based on observations of the development of *P. aeruginosa* and *S. aureus* biomass under mixed-species conditions across 8, 24 and 48 hours **(Figure 3)**, we measured aggregate sizes for *S. aureus* and biomass thickness for *P. aeruginosa* across subsequent time points **(Figures 4B and C)**. At 8 hours, *S. aureus* biomass consisted of a majority of aggregates (~94%) in the size range of <25 *μ*m^2^. At 24 and 48 hours, this proportion was observed to significantly decline, with ~80% of aggregates in the size range of <25 *μ*m^2^, and a shift towards larger aggregate sizes in the range of 25-50 μm^2^ and 50-100 *μ*m^2^ (~11-12%). Notably, there was no significant difference in the aggregate size distributions of *S. aureus* across 24 and 48 hours. For *P. aeruginosa*, across 8, 24 and 48 hours, the biomass grew into thick mats, measuring 56+3 *μ*m at 8 hours, and 46+1 *μ*m and 44±4 *μ*m at 24 and 48 hours respectively.

**Figure 4:**
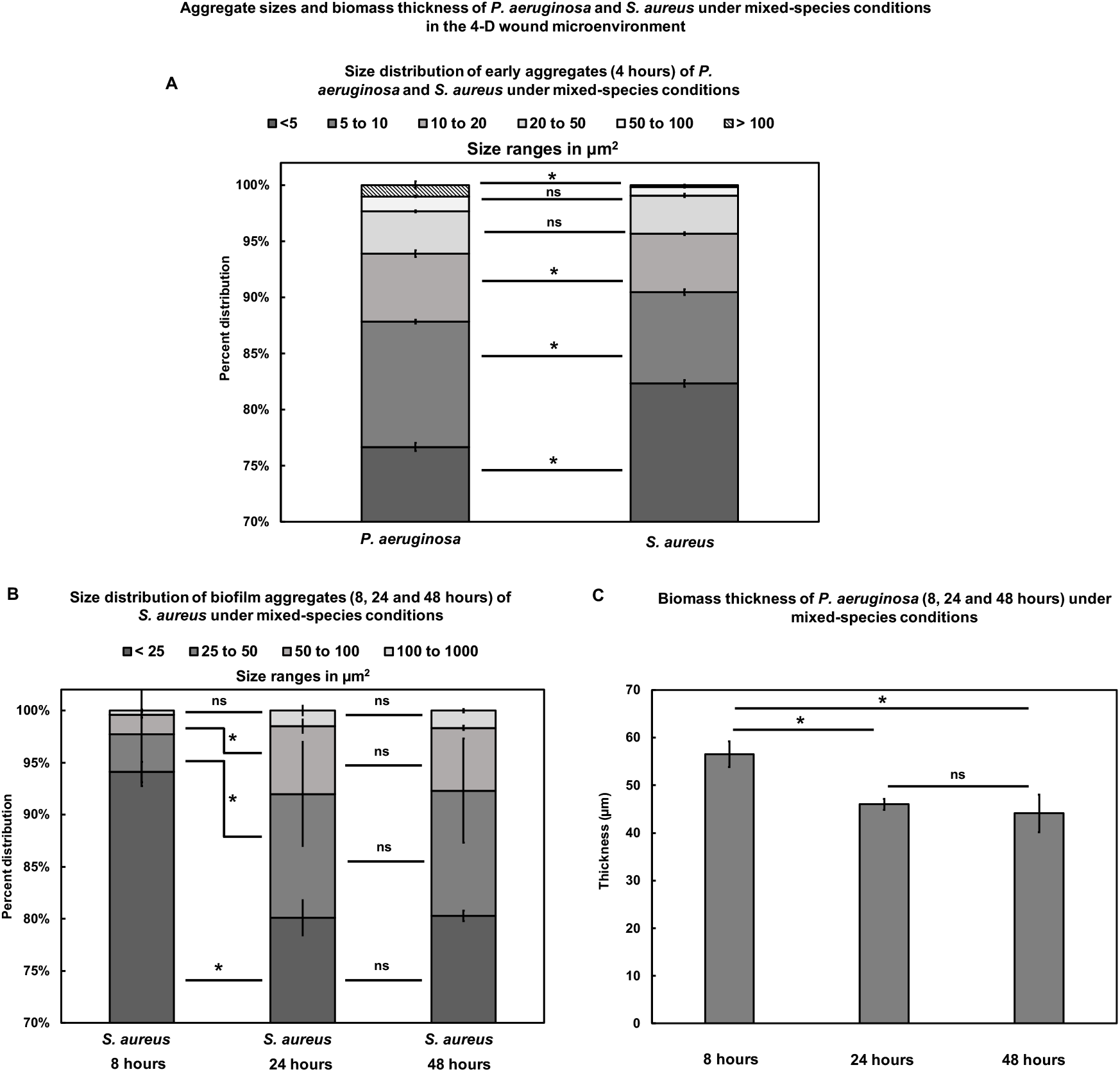
Aggregate size distribution and biomass thickness of *P. aeruginosa* and *S. aureus* under mixed-species conditions in the 4-D microenvironment. Aggregate sizes for *P. aeruginosa* and *S. aureus* in mixed-species biomass was measured at 4 hours, and for *S. aureus* across 8, 24 and 48 hours using the particle size analysis tool in ImageJ [47,48]. Biomass thickness for *P. aeruginosa* under mixed-species conditions across 8, 24 and 48 hours was measured using side view projection images of the Z-stacks in ImageJ [47,48]. **(A)** At 4 hours, the majority of *P. aeruginosa* and *S. aureus* biofilm aggregates under mixed-species conditions, consist of single bacterial cells or small aggregates (less than 5 *μ*m^2^). Notably, *S. aureus* consists of a larger percentage of small aggregates (less than 5 *μ*m^2^) as compared with *P. aeruginosa*. **(B)** Aggregate size analysis of *S. aureus* biomass across 8, 24 and 48 hours under mixed-species conditions, shows that the majority of *S. aureus* biomass consists of aggregates less than 25 *μ*m^2^ in size, with a significant shift towards larger aggregates seen at 24 and 48 hours (25-50 *μ*m^2^ and 50-100 *μ*m^2^). **(C)** Biomass thickness analysis of *P. aeruginosa* across 8, 24 and 48 hours under mixed-species conditions, showing *P. aeruginosa* growing to form thick biofilms. Error bars represent SEM, n=3, biological replicates; ns indicates no significant difference, a p-value ≤0.05 was considered significant (*).

Taken together, in the early mixed-species biomass, *P. aeruginosa* and *S. aureus* consist of discrete aggregates, with *P. aeruginosa* forming larger aggregates. Across subsequent growth, *S. aureus* retains its aggregate structure, albeit growing to form larger aggregates. On the other hand, *P. aeruginosa* grows into thick, mat-like biofilm structures, which appear to surround *S. aureus* aggregates.

### Co-localization of *P. aeruginosa* and *S. aureus* in mixed-species biofilms in the 4-D wound microenvironment

To determine the co-localization of *P. aeruginosa* and *S. aureus* in the mixed species biomass the 3D overlap analysis module of BiofilmQ v0.2.1 [41,42] was used. This tool measures the volumetric overlap fraction for each species, which is the volume of the species (channel) overlap in relation to the biomass of that species.

As seen in **Figure 5A**, at 4 hours, a very small fraction of the total biomass of *P. aeruginosa* was observed to overlap with *S. aureus*, with the non-overlapping fraction measuring ~99%. This could be due to the dispersed nature of the *P. aeruginosa* and *S. aureus* biomass, seen as discrete aggregates, under mixed-species conditions. The *P. aeruginosa* overlap fraction increases across the time points of 8, 24 and 48 hours, but continues to remain a very small fraction of the total *P. aeruginosa* biomass, accounting for ~2% of the species biomass at 48 hours. This indicates that a small proportion of *P. aeruginosa* overlaps with *S. aureus* in the mixed-species biomass, with the large majority of the *P. aeruginosa* biomass constituting the non-overlap fraction. This is possibly due to the growth of *P. aeruginosa* into thick, mat-like biofilms across time, resulting in small regions of overlap with *S. aureus* aggregates. On the other hand, for *S. aureus*, the biomass overlapping with *P. aeruginosa* at 4 hours constitutes ~28% of the total *S. aureus* biomass (**Figure 5B)**. The larger overlap fraction of *S. aureus* with *P. aeruginosa*, as compared with vice-versa, could be due to the larger proportion of smaller aggregates in the *S. aureus* in the early mixed-species biomass (**Figures 3A and 4A**). This could result in more *S. aureus* (given the smaller aggregate sizes) in proximate distance to P. aeruginosa, in contrast to larger aggregates of *P. aeruginosa*, with substantial biomass at the center of the aggregates. At 8 hours, the overlap fraction of *S. aureus* was observed to decrease to ~7%, followed by an increase across 24 and 48 hours (~30% and 28% respectively). This decrease in the *S. aureus* overlap fraction at 8 hours could result from a combination of several factors such as growth and density of aggregates of *S. aureus*, as well as increase in *P. aeruginosa* biomass.

**Figure 5:**
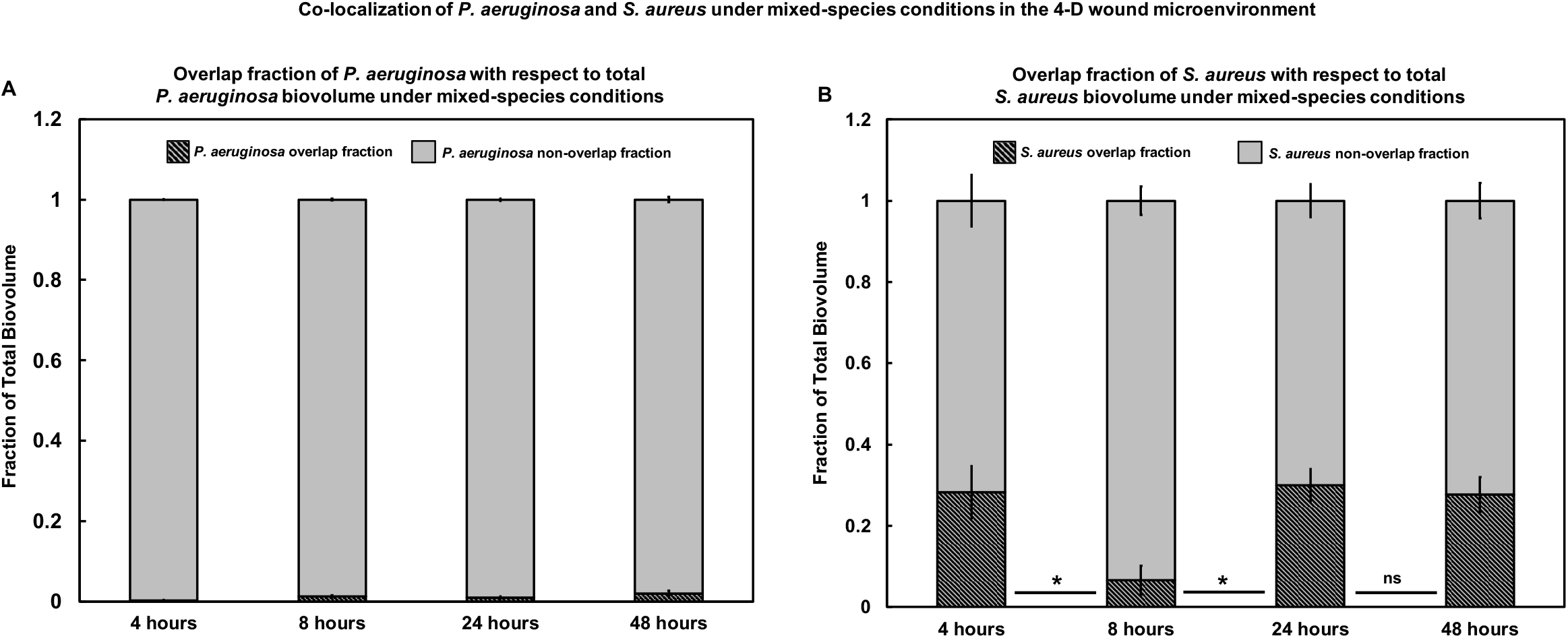
Co-localization of *P. aeruginosa* and *S. aureus* biomass under mixed-species conditions in the 4-D microenvironment. The relative organization of *P. aeruginosa* and *S. aureus* with respect to each other was measured using the 3D overlap parameter in BiofilmQ v0.2.1 [41,42]. **(A)** The overlap fraction of *P. aeruginosa* with respect to the total *P. aeruginosa* biovolume, shows a small fraction of *P. aeruginosa* overlapping with *S. aureus*, with the non-overlap fraction comprising the vast majority of the biomass. **(B)** The overlap fraction of *S. aureus* with respect to the total *S. aureus* biovolume, shows a larger proportion of *S. aureus* overlapping with *P. aeruginosa* (as compared with vice versa), with the overlap fraction decreasing at 8 hours, followed by an increase at 24 and 48 hours. Error bars represent SEM, n=3, biological replicates; ns indicates no significant difference, a p-value ≤0.05 was considered significant (*).

Taken together, it is evident that in the mixed-species biomass, *P. aeruginosa* and *S. aureus* exist in close proximity, and the differences in overlap fractions between the two species and across time points indicates distinct organization of the two species in the mixed-species biomass.

### Spatial organization of *P. aeruginosa* and *S. aureus* in mixed-species biofilms in the 4-D wound microenvironment

To study the spatial organization of *P. aeruginosa* and *S. aureus* across the mixed-species biomass, the MGV for each Z-slice in a given channel was calculated using the ‘measure stack’ tool in ImageJ [47,48]. MGVs were normalized to the highest MGV in a given dataset, and plotted as a distribution of biomass over Z-thickness.

As seen in **Figure 6A**, at 4 hours, the normalized MGV distribution of both *P. aeruginosa* and *S. aureus* in the mixed-species biomass is observed to coincide with the normalized MGV distribution of the host cell scaffold (the DAPI channel) at the lower end of the Z-stack. This indicates that in the 4-D microenvironment, early mixed-species biomass, including biofilm aggregates, form in close association with the host cell scaffold and IVWM, with a dispersion thickness (defined as total peak width) between 50-60 *μ*m for *P. aeruginosa*, 25-30 *μ*m for *S. aureus* and ~50 *μ*m for the DAPI channel. For both species, a proportion of biomass is also seen to be dispersed across the upper regions of the Z-stack (dispersion thickness 90-95 *μ*m), indicating the presence of free-floating aggregates or planktonic cells on top of the biomass. At 8 hours, the normalized MGV distribution **(Figure 6B)** shows the *P. aeruginosa* biomass to be distributed across an increased region (as compared with 4 hours) (dispersion thickness ~80 *μ*m), with the *S. aureus* biomass observed in the lower regions of the Z-stack (dispersion thickness of 30-40 *μ*m). In doing so, the *S. aureus* biomass is observed to lie within the *P. aeruginosa* biomass. Further, at 24 hours **(Figure 6C)**, peak of the normalized MGV distribution *P. aeruginosa* and *S. aureus* biomass is at ~55 *μ*m thickness indicating biomass accumulation at this height. However, similar to 8 hours biofilms, *P. aeruginosa* biomass is distributed across a wider thickness while *S. aureus* biomass is seen within the *P. aeruginosa* biomass. At 48 hours **(Figure 6D)**, while the *P. aeruginosa* biomass distribution further increases (dispersion thickness ~100-110 *μ*m), *S. aureus* remains in the lower regions of the biomass (dispersion thickness ~45-50 *μ*m).

**Figure 6:**
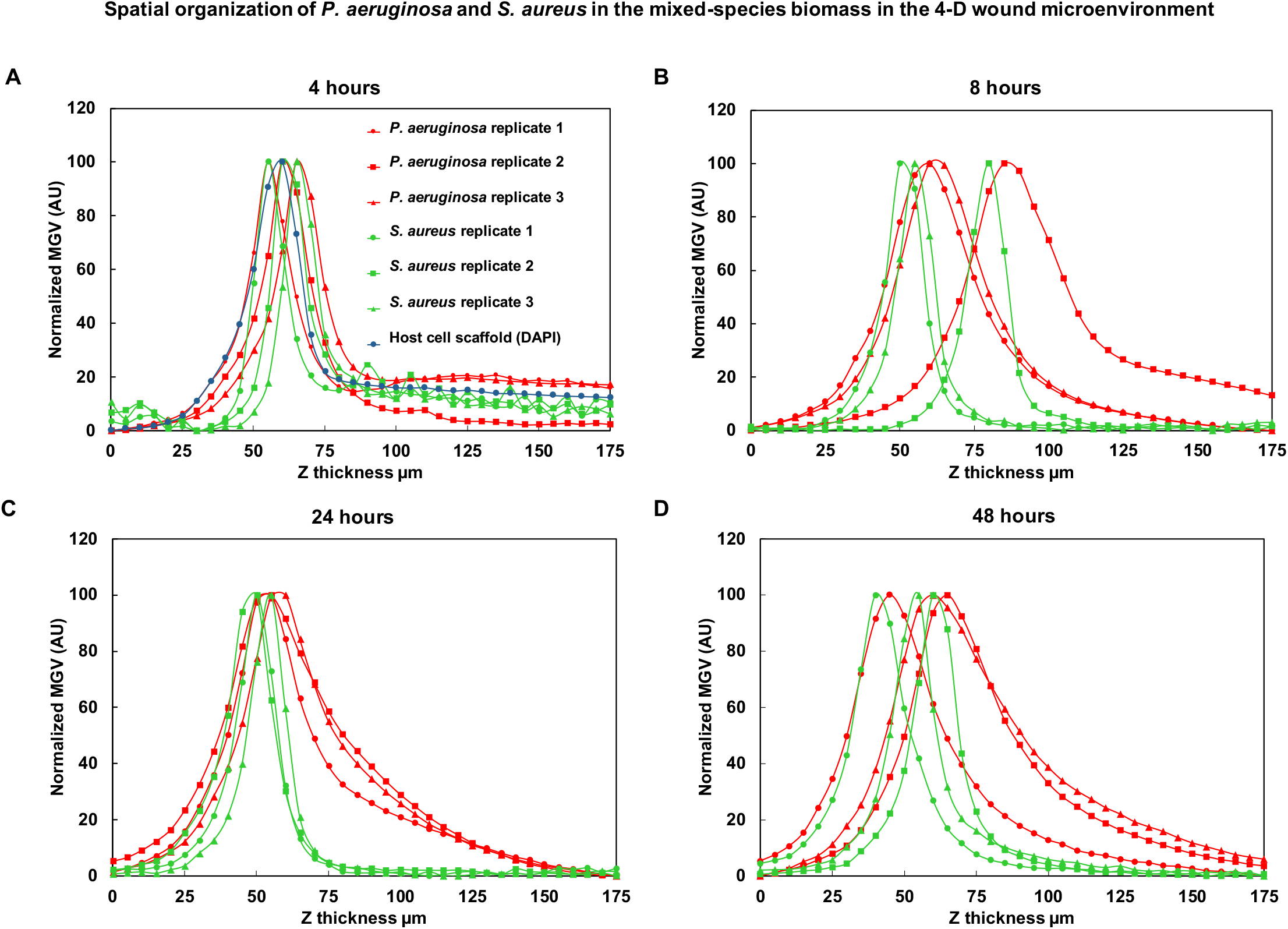
Spatial organization of *P. aeruginosa* and *S. aureus* under mixed-species conditions in the 4-D microenvironment. For *P. aeruginosa* and *S. aureus* in mixed-species biofilms across 4, 8, 24 and 48 hours in the 4-D microenvironment, the MGV for each Z-slice in a given channel was calculated using ImageJ [47,48]. MGVs were normalized to the highest MGV in a given dataset, and plotted as a distribution of biomass over Z-thickness. **(A)** At 4 hours, *P. aeruginosa* and *S. aureus* biomass is observed in proximity with the host cell scaffold (the DAPI channel), with *P. aeruginosa* and *S. aureus* distributed across the mixed-species biomass. **(B)** At 8 hours, *P. aeruginosa* biomass is distributed across a greater thickness in comparison to 4 hours, while the *S. aureus* continues to be seen at the lower end of the mixed-species biomass. **(C)** At 24 hours, *P. aeruginosa* biomass shows a wide distribution, with the *S. aureus* biomass observed to lie within the *P. aeruginosa* biomass. **(D)** At 48 hours, *P. aeruginosa* biomass shows a similar distribution to 24 hours, with *S. aureus* seen at the lower end of the mixed-species biomass. Each set of biological replicates for *P. aeruginosa* and *S. aureus* shown separately, n=3.

Taken together, in early mixed-species conditions in the 4-D microenvironment, the biomass of *P. aeruginosa* and *S. aureus* (likely aggregates and planktonic cells) are well-mixed, with the distribution coinciding with the presence of the host cell scaffold. Across subsequent time points, the growth of *P. aeruginosa* into mat-like biofilms, and the presence of *S. aureus* as aggregates, results in spatial reorganization of the biomass with *S. aureus* embedded in the lower regions of the *P. aeruginosa* biomass.

### Biomass composition of *P. aeruginosa* and *S. aureus* under mixed-species conditions in the 4-D wound microenvironment

Biomass for mixed-species *P. aeruginosa* and *S. aureus* biofilms across time points was measured using mean gray values (MGV) in ImageJ [47,48]. MGV represents the mean fluorescent intensity (sum of pixel values divided by number of pixels), and can therefore serve as a proxy for total biomass present in the visualized area. The MGV ratio of *P. aeruginosa* and *S. aureus* when grown together represents the relative composition of the two species in mixed-species biofilms, with a ratio of >1 representing more *P. aeruginosa* biomass and a ratio of <1 representing more *S. aureus* biomass.

As seen in **Figure 7A**, in the 4-D microenvironment, the ratio of the biomass of *P. aeruginosa* to *S. aureus* under mixed-species conditions increases from 4 hours to 24 hours, with similar values across 24 and 48 hours, indicating a higher relative biomass of *P. aeruginosa* across time points. At 4 hours, the ratio of the relative biomass of *P. aeruginosa* to *S. aureus* is 1.4±0.38, which indicates the nearly equal composition of both species in the mixed-species biofilm. At 24 hours, the ratio of the relative biomass of *P. aeruginosa* to *S. aureus* in the mixed-species biofilm is 2.62±0.19, which is nearly twice the ratio of the relative biomass at 4 hours **(Figure 7A)**. The ratio stabilizes across 24 and 48 hours, with the relative biomass ratio of *P. aeruginosa* to *S. aureus* at the final time point of 48 hours remaining nearly twice that at the earliest time point of 4 hours **(Figure 7A)**.

**Figure 7:**
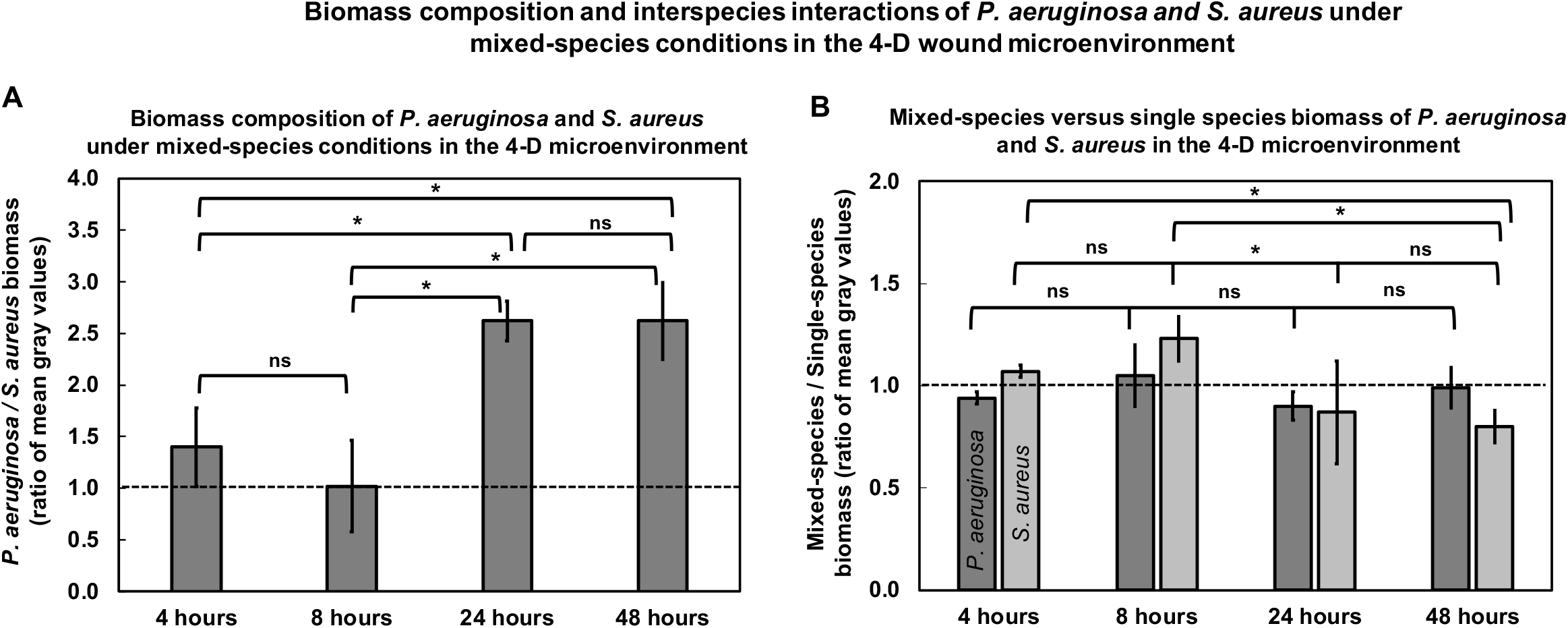
Biomass composition and interspecies interactions of *P. aeruginosa* and *S. aureus* under mixed-species conditions in the 4-D microenvironment. Biomass for *P. aeruginosa* and *S. aureus* biofilms, grown alone or together on HaCaT+HDFa scaffolds with IVWM across 4, 8, 24 and 48 hour time points was measured using mean gray values (MGV) in ImageJ [47,48]. **(A)** The ratio of *P. aeruginosa* to *S. aureus* in the mixed-species biomass shows the co-existence of both species, with a relative predominance of *P. aeruginosa* across time points. **(B)** The ratio of mixed- to single-species biomass for *P. aeruginosa* and *S. aureus* across 4, 8, 24 and 48 hour time points, indicates that in mixed-species biofilms, interspecies interactions could contribute to the impaired or inhibited growth of *S. aureus*. Error bars represent SEM, n=3, biological replicates; ns indicates no significant difference, a p-value ≤0.05 was considered significant (*).

Taken together, this indicates that the 4-D microenvironment supports the co-existence of *P. aeruginosa* and *S. aureus* in mixed-species biofilms, with *P. aeruginosa* steadily outcompeting *S. aureus*. This aligns with previous *in vivo* and clinical observations of wound biofilms [6,7,18,21,33,71], as well as biomimetic *in vitro* studies, where *P. aeruginosa* and *S. aureus* are seen to co-exist in wound-like conditions, with *P. aeruginosa* found in larger numbers [18,33]. The predominance of *P. aeruginosa* in the mixed-species biomass could be due to an inhibitory effect of *P. aeruginosa* on the growth of *S. aureus* [18,72,73] (widely observed under *in vitro* conditions), or due to the inherent growth dynamics of the individual bacterial species in the 4-D microenvironment, such as the increased growth of *P. aeruginosa* under mixed-species conditions [21].

### Possible interspecies interactions of *P. aeruginosa* and *S. aureus* under mixed-species conditions in the 4-D wound microenvironment

To parse out the possible roles of the inhibitory interspecies interactions and relative bacterial growth properties, we compared the MGV ratio for *P. aeruginosa* and *S. aureus* under mixed-species conditions to single-species conditions across time points, with a ratio of 1 representing equal biomass for the species across mixed- and single-species conditions, a ratio of >1 represents more biomass for the species under mixed-species conditions, and a ratio of <1 represents more biomass for the species under single-species conditions.

As seen in **Figure 7B**, the ratio of mixed- to single-species biomass for *P. aeruginosa* in the 4-D microenvironment remains close to 1, with no significant difference across time points. On the other hand, the ratio of mixed- to single-species biomass for *S. aureus* shows a significant decrease across 8 and 24 hours, followed by similar values across 24 and 48 hours. At 48 hours, the ratio of the relative biomass of S. aureus under mixed-species conditions as compared with single-species conditions is significantly less than the ratio of the relative biomass at 4 hours. This indicates that in mixed-species biofilms, the growth of *S. aureus* is impaired or inhibited in comparison with that under single-species conditions, with the impaired growth observed across 8 and 48 hours. This could be due to well-studied interspecies interactions such as the active killing or inhibition of the growth of *S. aureus* by a range of *P. aeruginosa* exoproducts [72,73], which correlates with the increased accumulation of *P. aeruginosa* across time. At early time points, the possible inhibitory effect on S. aureus can be observed in the aggregate size distribution of S. aureus under single-species conditions when compared with mixed-species biomass **(Supplementary Figure 3)**. At 4 and 8 hours, *S. aureus* under single-species conditions showed a significant shift towards larger aggregate sizes, as compared with that observed under mixed-species conditions **(Supplementary Table 1)**. At 4 hours, this was seen as an increase in aggregates in the 10-20 *μ*m^2^ and 20-50 *μ*m^2^ size ranges, and at 8 hours this was seen as an increase in aggregates in the 25-50 *μ*m^2^ and 50-100 *μ*m^2^ size ranges **(Supplementary Figure 3A and B)**. On the other hand, *P. aeruginosa* biomass under single-species conditions grow into several micron-thick mats, similar to that under mixed-species conditions **(Supplementary Figure 3C)**.

At later time points, the biomass accumulation (notably that of *P. aeruginosa*) disrupts the host cell scaffold, seen as diffusion of the blue nuclear stain at 24 and 48 hours **(Figure 3A and Supplementary Figure 1)**. This partial loss of scaffold structure overlaps with the significant decline in *S. aureus* biomass, observed across 8 and 48 hour time points **(Figure 7B)**. This could indicate that the presence of the host cell scaffold affords some protection of *S. aureus* biofilms from the possible inhibitory effects of *P. aeruginosa*. Host cell scaffold-associated *S. aureus*, as adherent biomass or biofilm aggregates [28,68,74,75], could possess increased tolerance or growth advantages, including protection against virulence factors and exoproducts.

### Proposed fine-tuned model of the structure and organization of mixed-species *P. aeruginosa* and *S. aureus* biomass in the wound bed

Taken together, the composite 4-D microenvironment consisting of co-cultured (fixed) HaCaT+HDFa cell scaffolds and the IVWM, supports the co-existence of *P. aeruginosa* and *S. aureus* under mixed-species conditions with characteristic features relevant to biofilms in the wound bed. In early mixed-species conditions, *P. aeruginosa* and *S. aureus* are seen as biofilm aggregates or planktonic cells randomly distributed across the biomass, and in close association with the host cell scaffold. At this stage, *P. aeruginosa* is seen to form larger aggregates as compared with *S. aureus*, which could possibly explain the differences in overlap fraction between the biomass of the two species. Further, based on biomass composition, the early mixed-species biomass shows the nearly equal presence of both *P. aeruginosa* and *S. aureus*. In subsequent growth of the mixed-species biomass, *S. aureus* was observed to retain its aggregate structure, albeit forming larger aggregates. On the other hand, *P. aeruginosa* grew into large, dense mat-like structures. The subsequent growth of mixed-species biomass was associated with distinct changes in co-localization and spatial organization across the two species. Notably, *S. aureus* aggregates were seen embedded in the lower parts of, and surrounded by the, *P. aeruginosa* biomass. Further, the biomass composition of the mixed-species biomass revealed a predominance of *P. aeruginosa* over time, likely a result of interspecies interactions resulting in *P. aeruginosa* inhibiting the growth of *S. aureus* [72,73].

In chronic wound biofilms, *P. aeruginosa* and *S. aureus* show distinct spatial segregation, with *S. aureus* seen as smaller aggregates closer to the wound surface [6], and *P. aeruginosa* found as larger aggregates in the deeper regions of the wound. This has been partially attributed to the virulence features of *P. aeruginosa*, including motility, the ability to destroy polymorphonuclear leukocytes, adapt to low-oxygen [76,77] conditions and produce tissue remodeling enzymes [76–80]. It is important to note that this non-random distribution occurs over prolonged periods of non-healing, and will therefore be influenced by immune factors and tissue remodeling. Given this, it is very likely that in the initial phases of bacterial colonization, *P. aeruginosa* and *S. aureus* coexist in close proximity [21]. This could result from colonization with one pathogen predisposing establishment of the second pathogen, or infection with both pathogens could occur over short intervals of time. Regardless, the co-existence of both pathogens in the wound bed would result in a range of interactions, likely to influence the subsequent structure and organization of mixed-species biofilms in the wound bed.

In this study, the composite 4-D wound microenvironment with co-cultured (fixed) HaCaT+HDFa scaffolds and IVWM recapitulates the co-existence and interactions of *P. aeruginosa* and *S. aureus* in the presence of host and matrix elements **(Figure 8)**. Based on our findings, under these wound-like conditions, *P. aeruginosa* and *S. aureus* grow to form mixed-species biomass with distinct structure and organization, resulting in smaller aggregates of *S. aureus* embedded in large, dense, mat-like *P. aeruginosa* biofilms. Over time, the combined effects of bacterial virulence factors and tissue remodeling enzymes (not recapitulated in the 4-D microenvironment), particularly that of *P. aeruginosa* [78–80], could result in *P. aeruginosa* migrating to the deeper regions of the wound, while *S. aureus* remains as smaller aggregates closer to the wound surface. This would result in niche-partitioning, such as that seen in chronic, non-healing wounds, where *P. aeruginosa* and *S. aureus* biofilm aggregates are distributed across different regions.

**Figure 8:**
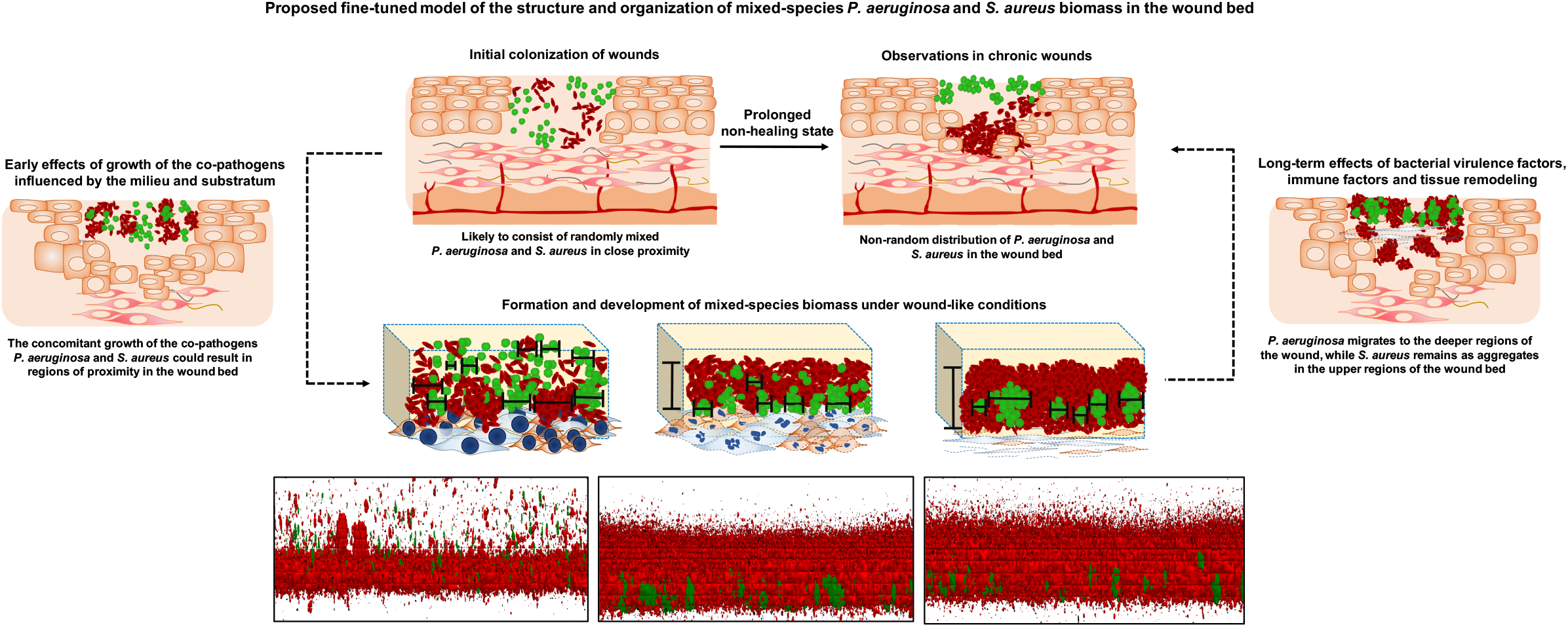
Proposed fine-tuned model of the structure and organization of mixed-species *P. aeruginosa* and *S. aureus* biofilms in the wound bed. During initial colonization and growth in the wound bed, *P. aeruginosa* and *S. aureus* are likely to exist in regions of close proximity. In this study, the composite 4-D microenvironment consisting of co-cultured (fixed) HaCaT+HDFa scaffolds with IVWM recapitulates the concomitant and proximate presence of *P. aeruginosa* and *S. aureus* in the wound bed. Based on our findings, under these wound-like conditions, *P. aeruginosa* and *S. aureus* grow to form mixed-species biomass with distinct structure and organization, resulting in smaller aggregates of *S. aureus* embedded in large, dense, mat-like *P. aeruginosa* biofilms. Further, the biomass composition revealed a predominance of *P. aeruginosa* over time, likely a result of interspecies interactions resulting in *P. aeruginosa* inhibiting the growth of *S. aureus*. Over time, the combined effects of bacterial virulence factors, immune factors and tissue remodeling could result in *P. aeruginosa* migrating to the deeper regions of the wound, while *S. aureus* remains as smaller aggregates closer to the wound surface. This could explain observations in chronic, non-healing wounds, where *P. aeruginosa* and *S. aureus* biofilm are distributed across distinct compartments.

It is important to consider the fact that the 4-D microenvironment is not inclusive of all factors relevant to the wound infection state, most notably the absence of live host cells and immune factors. Given this the insights into structure and organization of mixed-species biofilms are based on the influence of the biomimetic host substratum, and matrix and biochemical factors.

## CONCLUSIONS

The composite 4-D microenvironment recapitulates key features of the wound infection state, such as a reconstructed host cell surface and wound milieu. We leveraged this recapitulated microenvironment to characterize structure of mixed-species *P. aeruginosa* and *S. aureus* biofilms across multiple levels of organization, such as aggregate dimensions and biomass thickness, species co-localization and organization within the biomass, biomass composition and interspecies interactions. Mixed species biomass of *P. aeruginosa* and *S. aureus* grown in the recapitulated microenvironment, display distinct structure and organization, that varies between species and across biomass growth. Overall, several features of the mixed-species biomass in the 4-D microenvironment align with clinical and *in vivo* observations of wound biofilms [7–9], such as the formation of biofilm aggregates, coexistence of both pathogens with a predominance of *P. aeruginosa*, and indication of possible inhibitory effects of *P. aeruginosa* on *S. aureus*. Given this, the platform provides insights into mixed-species biomass features when *P. aeruginosa* and *S. aureus* are in close proximity, which can be placed in the context of the current understanding of *P. aeruginosa* and *S. aureus* organization in the wound bed.

While the fixed co-cultured host cell scaffold provides a biomimetic substratum, the wound bed consists of proliferating and migrating keratinocytes and fibroblasts, which are likely to influence host-biofilm interactions. Given that maintaining host cell viability in the presence of bacterial growth poses technical challenges [53], fixed host cell scaffolds provide a semi-inert, 3-D biomimetic substratum for biofilm growth, and DAPI-stained nuclei indicate the presence of host cells in relation to the mixed-species biomass. Nevertheless, future modifications of the platform could be optimized to allow for the presence of live host cells that would more closely mimic the host cell-biofilm interface. This would be particularly relevant in the context of studying initial colonization and bacterial adherence in the wound bed. Another limitation of this study is the absence of differentiation of viable and non-viable mixed-species biomass in the 4-D microenvironment. For this, optimizing the platform for live-dead staining would provide additional insights into the metabolic structure of the mixed-species biomass [50], which would be interesting given the chemical composition of the IVWM. In addition to probing mixed-species biofilm structure and organization, the 4-D microenvironment could also be developed into a platform for evaluating anti-biofilm approaches, under wound-relevant conditions. For this, in addition to structural and organizational features, optimizing parameters such as the assessment of viability and metabolic activity of biofilm cells would be useful. This could provide high-throughput screening of potential anti-biofilm treatments, with structural and functional readouts.

Taken together, the 4-D wound microenvironment represents a composite *in vitro* model that can be leveraged for a range of studies, including the formation and development of wound biofilms, and the evaluation of novel treatment approaches. For this, the reconstructed nature of the 4-D platform lends itself well not only for multi-level and multi-parameter insights, but also for parsing the roles of various host, matrix, biochemical, and microbial factors, including the interplays between them.

## Funding

KSK’s appointment is supported by the Ramalingaswami Re-entry Fellowship (BT/HRD/16/2006), Department of Biotechnology, Government of India. RD’s appointment is supported by the Har Gobind Khorana-Innovative Young Biotechnologist Award (HGK-IYBA), Department of Biotechnology, Government of India to KSK (BT/12/IYBA/2019/05). This study is funded by the Har Gobind Khorana-Innovative Young Biotechnologist Award (HGK-IYBA), Department of Biotechnology, Government of India to KSK (BT/12/IYBA/2019/05).

## Data availability statement

Data generated in this study will be made available by the authors upon request.

## Declaration of competing interest

The authors declare that they have no known competing financial interests or personal relationships that could have appeared to influence the work reported in this paper.

## CRediT authorship contribution statement

**Radhika Dhekane:** Methodology, Investigation, Validation, Writing – original draft. **Shreeya Mhade:** Methodology, Investigation, Validation, Writing – original draft. **Karishma S. Kaushik:** Conceptualization, Methodology, Formal analysis, Project administration, Supervision, Writing – original draft, Writing – review & editing, Funding acquisition.

## Author Contributions

RD performed the experimental work, did parts of the image analysis and wrote parts of the first draft of the manuscript. SM did parts of the image analysis and wrote parts of the first draft of the manuscript. KSK developed the research idea, supervised the work, wrote the first draft of the manuscript, and edited the final draft of the manuscript.

## Acknowledgements

We thank M Vandana and Sujaya Ingale, NCL-Innovation Park, Pune, India for technical assistance with confocal microscopy.

## SUPPLEMENTARY FIGURE LEGENDS

**Supplementary Figure 1: Co-cultured HaCaT+HDFa scaffold changes in the presence of mixed-species biomass over time.** The presence of mixed-species biomass results in progressive destruction of the host cells in the co-cultured (fixed) HaCaT+HDFa scaffold over time. This is observed as loss of nuclei architecture and diffusion of the blue nuclear stain across 24-48 hours. The red and green fluorescence channels corresponding to *P. aeruginosa* and *S. aureus* biomass are not shown.

**Supplementary Figure 2: Single-species biomass of *P. aeruginosa* and *S. aureus* biofilms in the 4-D microenvironment.** *P. aeruginosa* (PAO1-pUCP18, mCherry) and *S. aureus* (Strain AH 133-pAH13, GFP) were seeded separately in IVWM on fixed HaCaT+HDFa scaffolds. Following incubation at 37°C, biomass was imaged at 4, 8, 24 and 48 hours. **(A)** Representative images of single-species *P. aeruginosa* biomass in the 4-D microenvironment across time points. The blue stain represents the DAPI-stained host cell nuclei; the diffusion of the nuclear stain across subsequent time points is likely due to destruction of host cell structure with bacterial growth. **(B)** Representative images of single-species *S. aureus* biomass in the 4-D microenvironment across time points.

**Supplementary Figure 3: Aggregate size distribution and biomass thickness of *P. aeruginosa* and *S. aureus* under single-species conditions in the 4-D microenvironment.** Aggregate sizes for *P. aeruginosa* and *S. aureus* in single-species biomass was measured at 4 hours, and for *S. aureus* across 8, 24 and 48 hours using the particle size analysis tool in ImageJ [47,48]. Biomass thickness for *P. aeruginosa* across 8, 24 and 48 hours was measured using side view projection images of the Z-stacks in ImageJ [47,48]. **(A)** At 4 hours, the majority of *P. aeruginosa* and *S. aureus* biofilm aggregates under single-species conditions, consisted of single bacterial cells or small aggregates (less than 5 *μ*m^2^), with no significant difference across aggregate size distributions. **(B)** Aggregate size analysis of *S. aureus* biomass across 8, 24 and 48 hours under single-species conditions, showing the majority of *S. aureus* biomass as aggregates less than 25 *μ*m^2^ in size, followed by 25-50 *μ*m^2^. There is a significant increase in the percentage of aggregates less than 25 *μ*m^2^ in size at 24 hours. **(C)** Biomass thickness analysis of *P. aeruginosa* across 8, 24 and 48 hours under single-species conditions, showing *P. aeruginosa* growing to form thick biofilms. Error bars represent SEM, n=3, biological replicates; ns indicates no significant difference, a p-value ≤0.05 was considered significant (*).

## SUPPLEMENTARY TABLES

**Supplementary Table 1:**
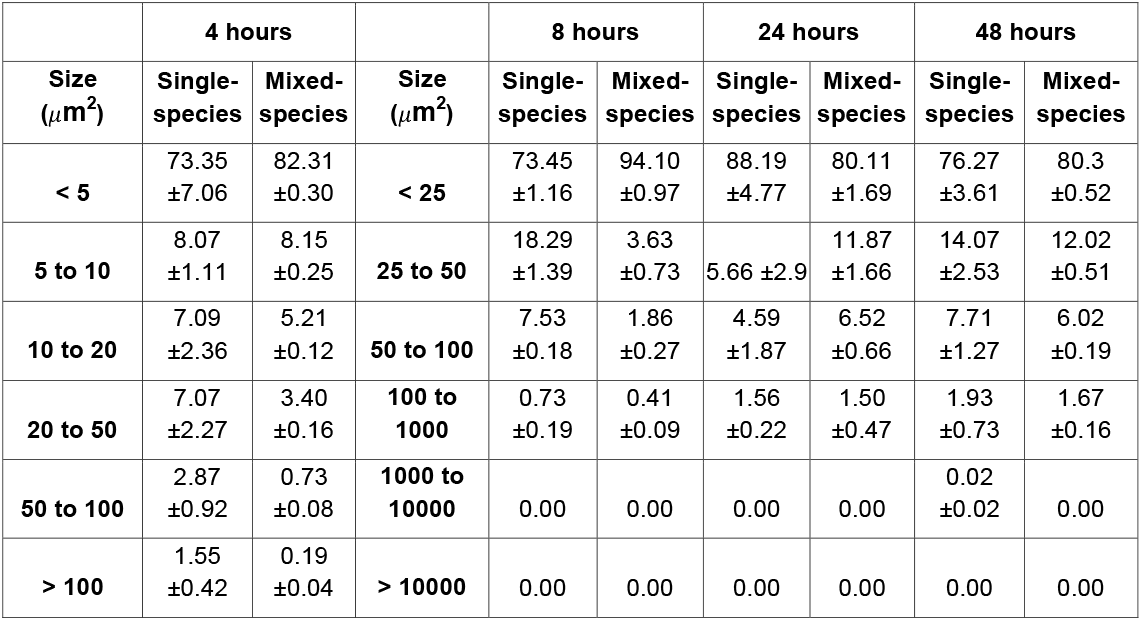
Percentage of the aggregate size distribution of *S. aureus* under single-species and mixed-species conditions over time in the 4-D microenvironment. (Error represents SEM, n=3, biological replicates).

## Notes

### Competing Interest Statement

The authors have declared no competing interest.

