## Supplementary material for "Adding a new dimension: Multi-level structure and organization of mixed-species *Pseudomonas aeruginosa* and *Staphylococcus aureus* biofilms in a 4-D wound microenvironment": Suppl Figure 1

### Changes in HaCaT + HDFa scaffold structure in the presence of mixed-species biofilms over time

#### 4 hours

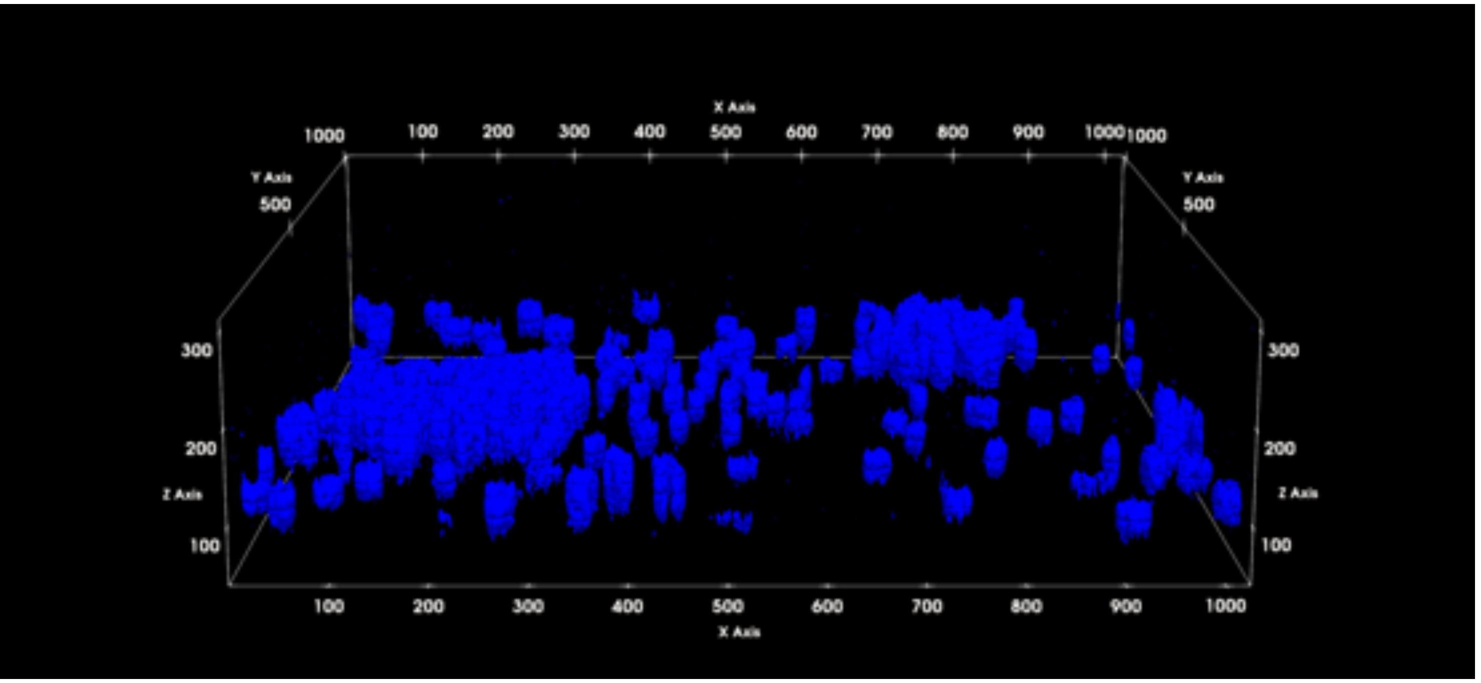


#### 8 hours

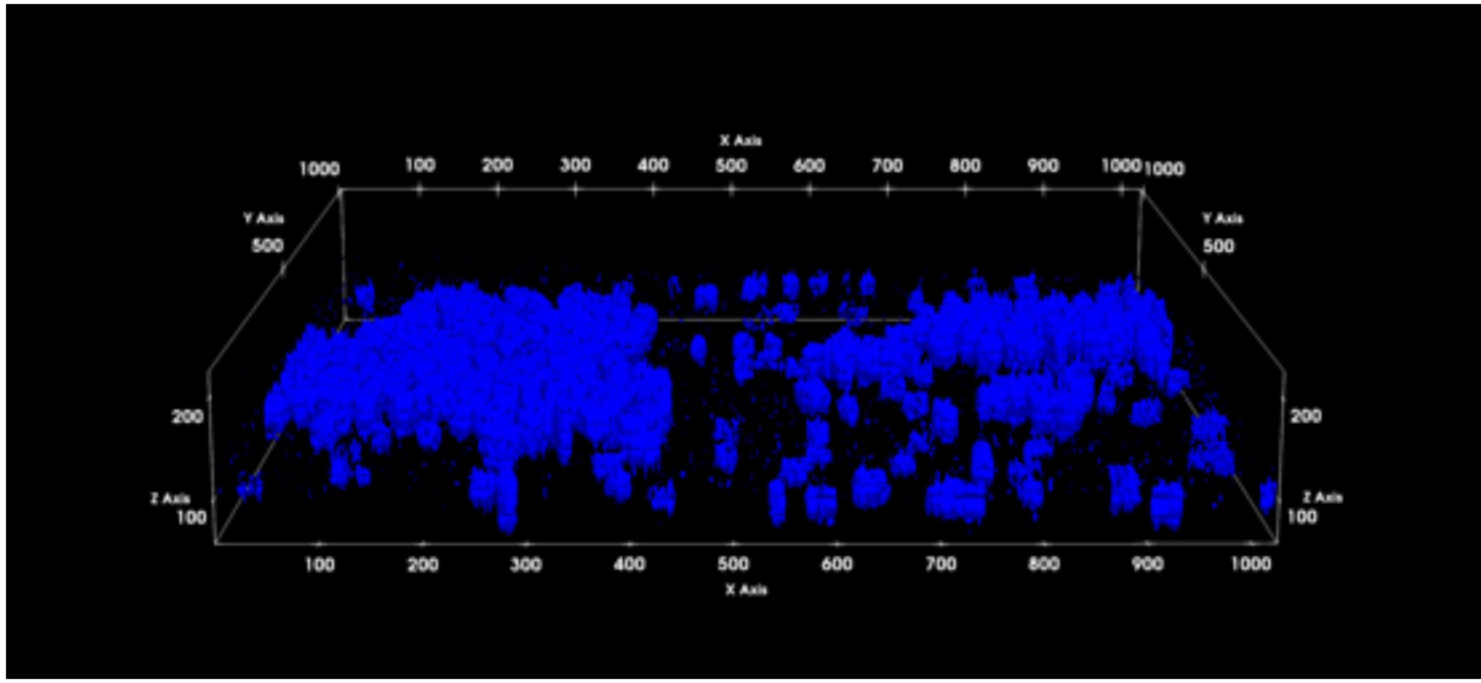


#### 24 hours

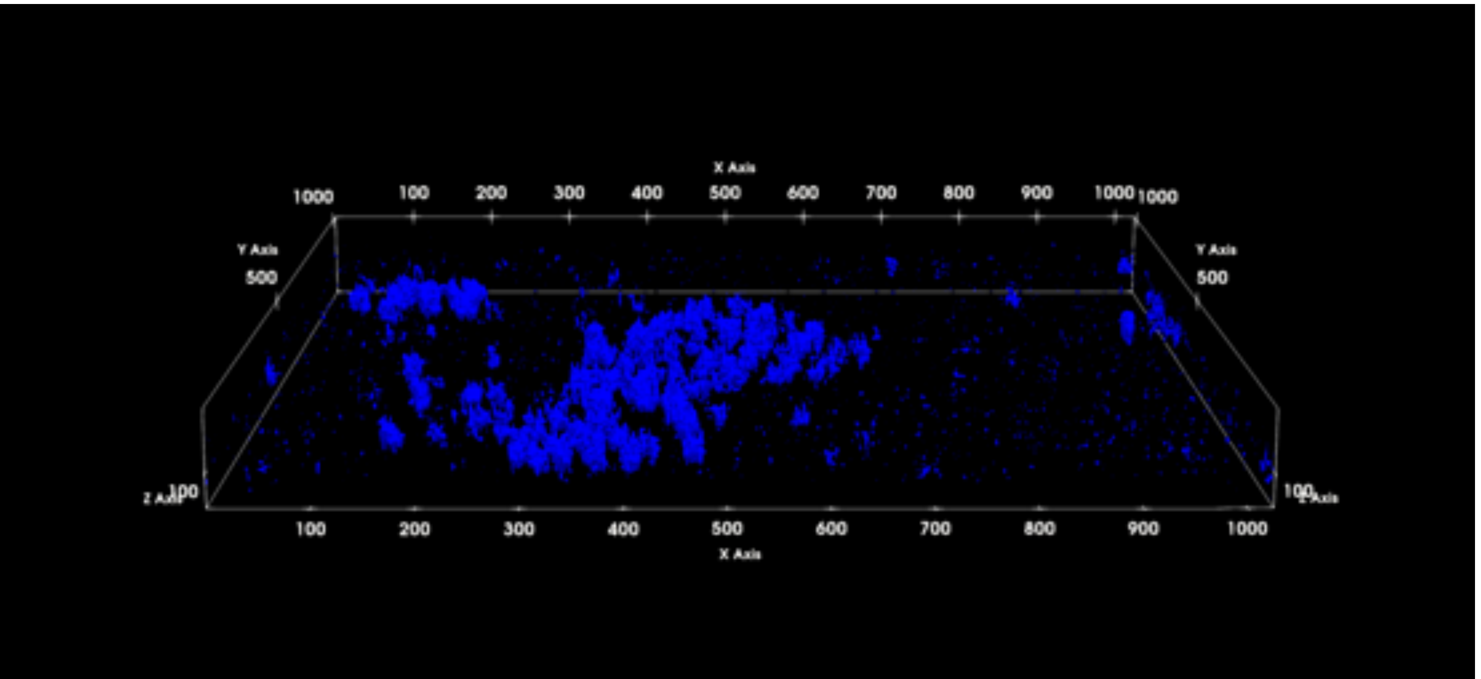


#### 48 hours

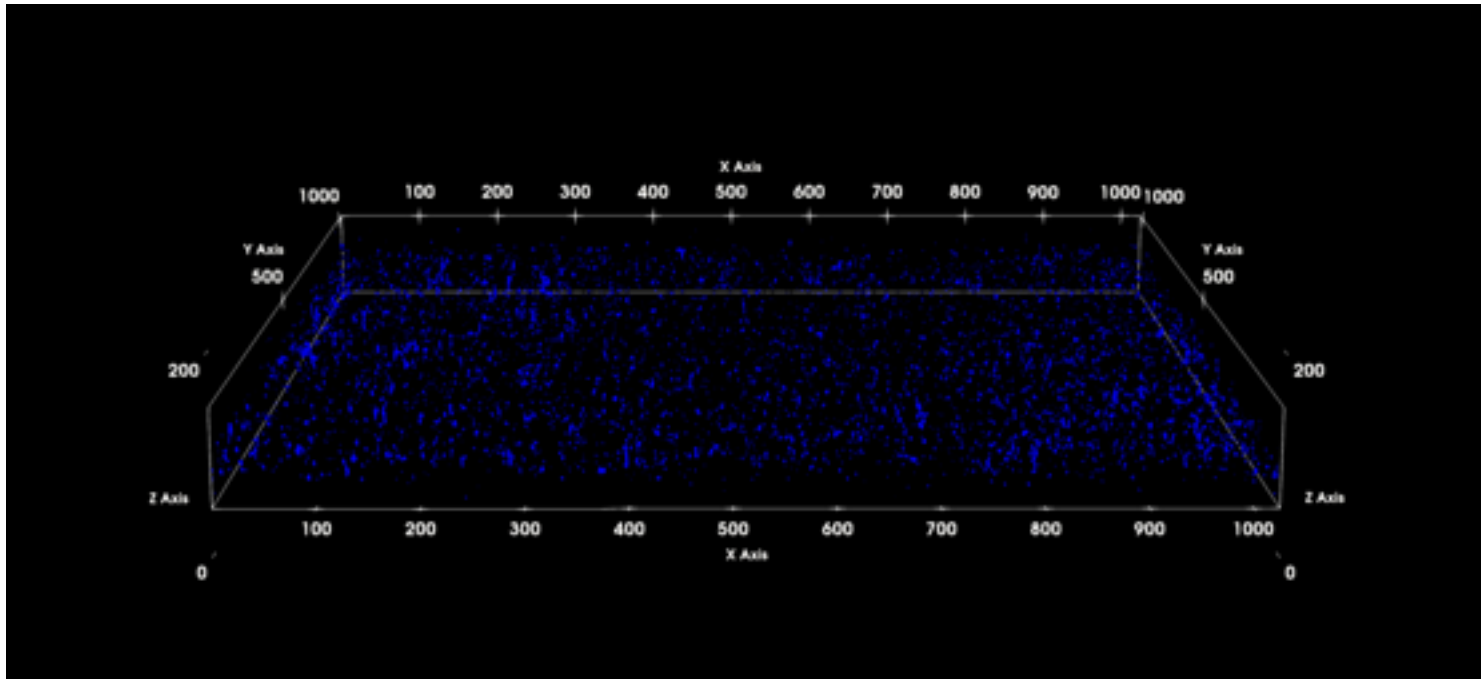
