## Supplementary material for "Adding a new dimension: Multi-level structure and organization of mixed-species *Pseudomonas aeruginosa* and *Staphylococcus aureus* biofilms in a 4-D wound microenvironment": Suppl Figure 2

**Single-species *P. aeruginosa* and *S. aureus* biofilms in the composite 4-D microenvironment**
**HaCaT + HDFa scaffolds with IVWM**

**A**

**Single-species *P. aeruginosa* biofilms**

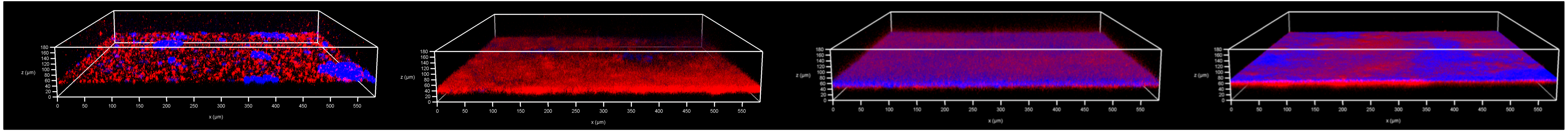

**4 hours** | **8 hours** | **24 hours** | **48 hours**

**B**

**Single-species *S. aureus* biofilms**

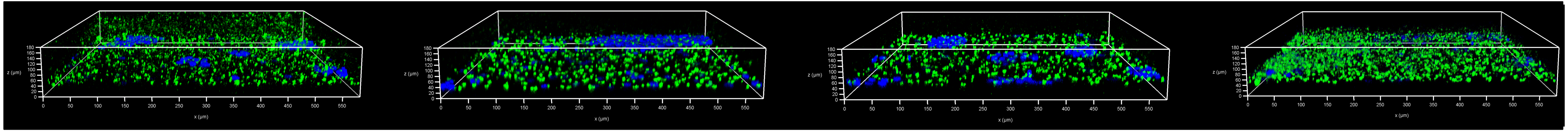

**4 hours** | **8 hours** | **24 hours** | **48 hours**
