## Supplementary material for "Adding a new dimension: Multi-level structure and organization of mixed-species *Pseudomonas aeruginosa* and *Staphylococcus aureus* biofilms in a 4-D wound microenvironment": Suppl Figure 3

**Aggregate sizes of *P. aeruginosa* and *S. aureus* under single-species conditions in the 4-D wound microenvironment**

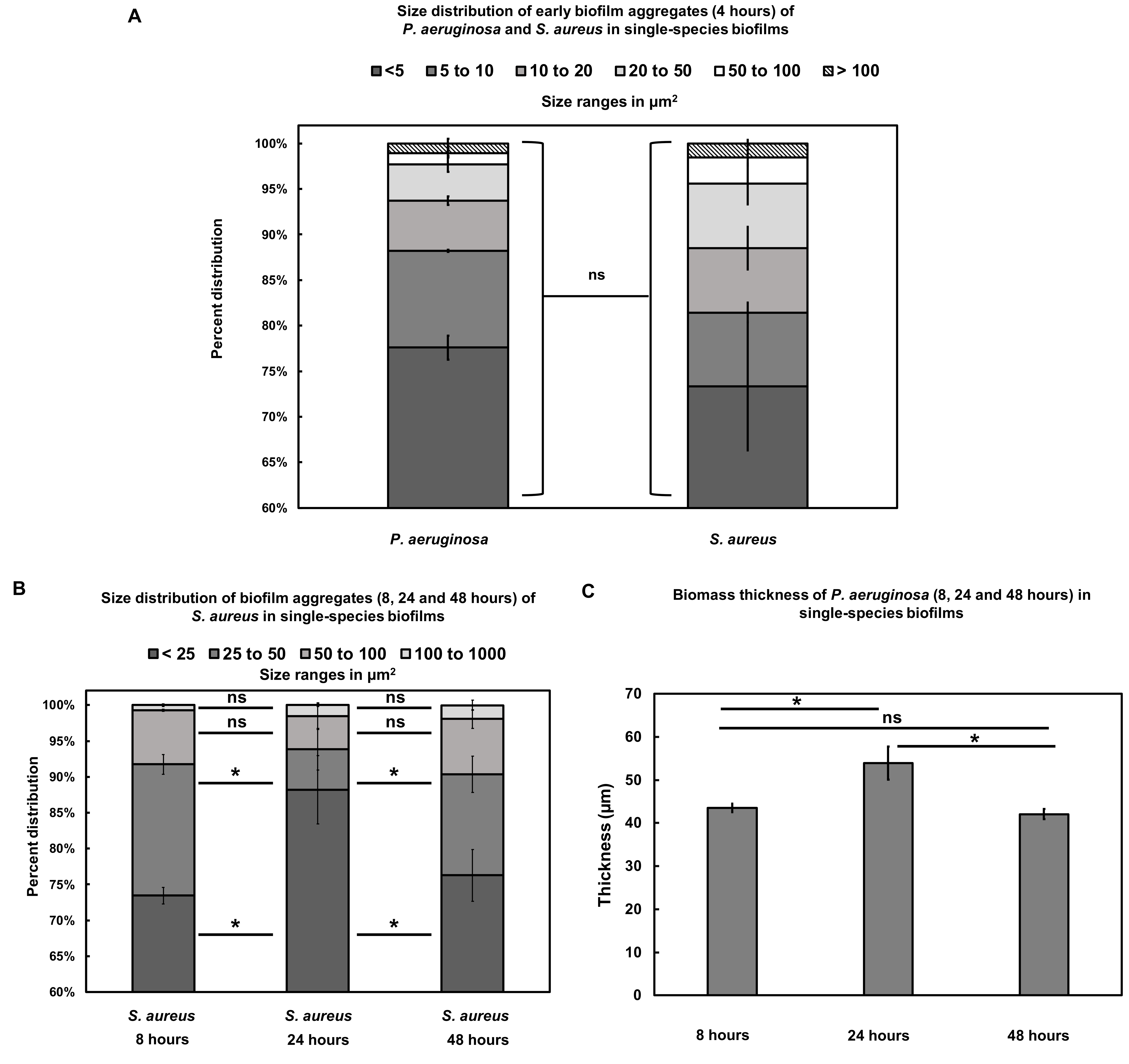
